# grandR: a comprehensive package for nucleotide conversion sequencing data analysis

**DOI:** 10.1101/2022.09.12.507665

**Authors:** Teresa Rummel, Lygeri Sakellaridi, Florian Erhard

## Abstract

Metabolic labeling of RNA is a powerful technique for studying the temporal dynamics of gene expression. Nucleotide conversion approaches greatly facilitate the generation of data but introduce challenges for their analysis. We here present grandR, a comprehensive package for quality control, differential gene expression analysis, kinetic modeling, and visualization of such data. We compare several existing methods for inference of RNA synthesis rates and half-lives using progressive labeling time courses. We demonstrate the need for recalibration of effective labeling times and introduce a Bayesian approach to study the temporal dynamics of RNA using snapshot experiments.

## Background

The RNA expression level of a gene is governed by the interplay of RNA synthesis and degradation. While RNA-seq can easily obtain transcriptome-wide snapshots of gene expression profiles in a single experiment, it remained difficult to directly measure the temporal dynamics of gene regulation as consequences of changes in the rates of RNA synthesis and degradation, e.g. due to external stimuli.

To overcome this limitation, techniques involving metabolic labeling of RNA have been developed. Metabolic RNA labeling uses 4-thiouridine (4sU) or other nucleoside analogs, which are introduced into living cells and incorporated into nascent RNA. First approaches physically separated labeled and unlabeled RNA by thiol-specific biotinylation and affinity purification and sequenced separate libraries of these fractions. This approach has been used to study RNA processing [1], transient RNA expression [2], kinetics of RNA polymerases [3] or the dynamics of RNA expression [4]. Existing protocols for purification of labeled RNA are highly laborious and require substantial amounts of RNA. In addition, contamination with background total RNA in the labeled RNA fraction must be controlled for [4], and normalization is challenging [5].

Recently, several approaches have been proposed that circumvent the purification [6–8]: Before sequencing, RNA is treated with chemical agents to specifically convert 4sU into cytosines or cytosine analogs. Thus, labeled and unlabeled RNAs can in principle be differentiated based on T-to-C mismatches in sequencing reads without the need to physically purify labeled RNA. A major advantage of the nucleotide conversion approach, aside from lower requirements of starting material and a simplified experimental workflow, is that it can be combined with more specialized protocols, e.g. to profile transcription start sides [9] or ribosome occupancy [10]. Furthermore, we and others have combined 4sU conversion with single cell RNA-seq to study the heterogeneity of gene regulation [11–13].

A limitation of 4sU conversion approaches is that concentrations of 4sU that are tolerated by cells commonly only replace 1 in 40 uridines by 4sU. Thus, a considerable number of reads originating from labeled RNA does not cover any 4sU incorporation site. The percentage of such reads is in the order of 20-80% and depends on the ratio of 4sU and normal uridine available for incorporation into nascent RNA, the read length, and the uridine content of RNA. The pioneering studies employing 4sU conversion exclusively focused on T-to-C reads. Despite underestimating labeled RNA, T-to-C reads alone can be used to estimate unbiased RNA half-lives in pulse-chase experiments [6] and to detect rapid changes of transcription upon drug treatment or acute depletion of transcription factors [14].

We previously proposed a statistical solution to quantify labeled and unlabeled RNA without bias due to limited 4sU incorporation: By using a Binomial mixture model, our GRAND-SLAM approach provides unbiased estimates of the percentage of labeled RNA per gene and its posterior distribution [15]. The posterior represents uncertainties in the quantification, mainly due to the scarcity of 4sU incorporation events, and has so far been used to filter out genes with inaccurate quantification [11]. Importantly, however, our Bayesian framework in principle allows to take these uncertainties further to downstream analyses such as estimation of RNA kinetics or gene expression changes.

Here, we present grandR, an R package to facilitate analyses of nucleotide conversion sequencing experiments. It includes new methods for quality control and recalibrating labeling times. grandR implements several methods to estimate RNA synthesis and degradation rates from progressive labeling experiments that have been applied previously by us and others [3,15–18]. Here, we compare these methods and show that the most accurate results are obtained by directly utilizing the posteriors from GRAND-SLAM to estimate the kinetic model. Furthermore, we propose a Bayesian hierarchical model to dissect the mode of gene regulation from snapshot experiments. To facilitate collaborative work and exploratory data analysis, grandR provides a comprehensive web-based data visualization and exploration tool.

## Results

### grandR overview

grandR is designed as a comprehensive and easy-to-use toolkit for all types of nucleotide conversion sequencing data such as SLAM-seq [6], Timelapse-seq [7] or TUC-seq [8]. Raw data is pre-processed, e.g. by our GRAND-SLAM tool [15]. Analysis workflows in grandR consist of high-level commands to (i) load data, (ii) filter genes according to user defined criteria, (iii) quality control, (iv) normalization and (v) kinetic modeling or differential gene expression analysis (Fig. 1A). Our GRAND-SLAM – grandR workflow advocates using systematic sample names to encode all metadata (Fig. 1B). Various experimental conditions that impact on the analysis strategy are accommodated by using specific parameters to grandR functions (Fig. 1C). grandR provides tools for visualizations of individual genes and summaries of a data set, which can be used programmatically or via a shiny-based web interface (Fig. 1D). All figures here were generated using grandR, and R notebooks to reproduce all analyses are provided as Supplementary data file 1.

**Figure 1:**
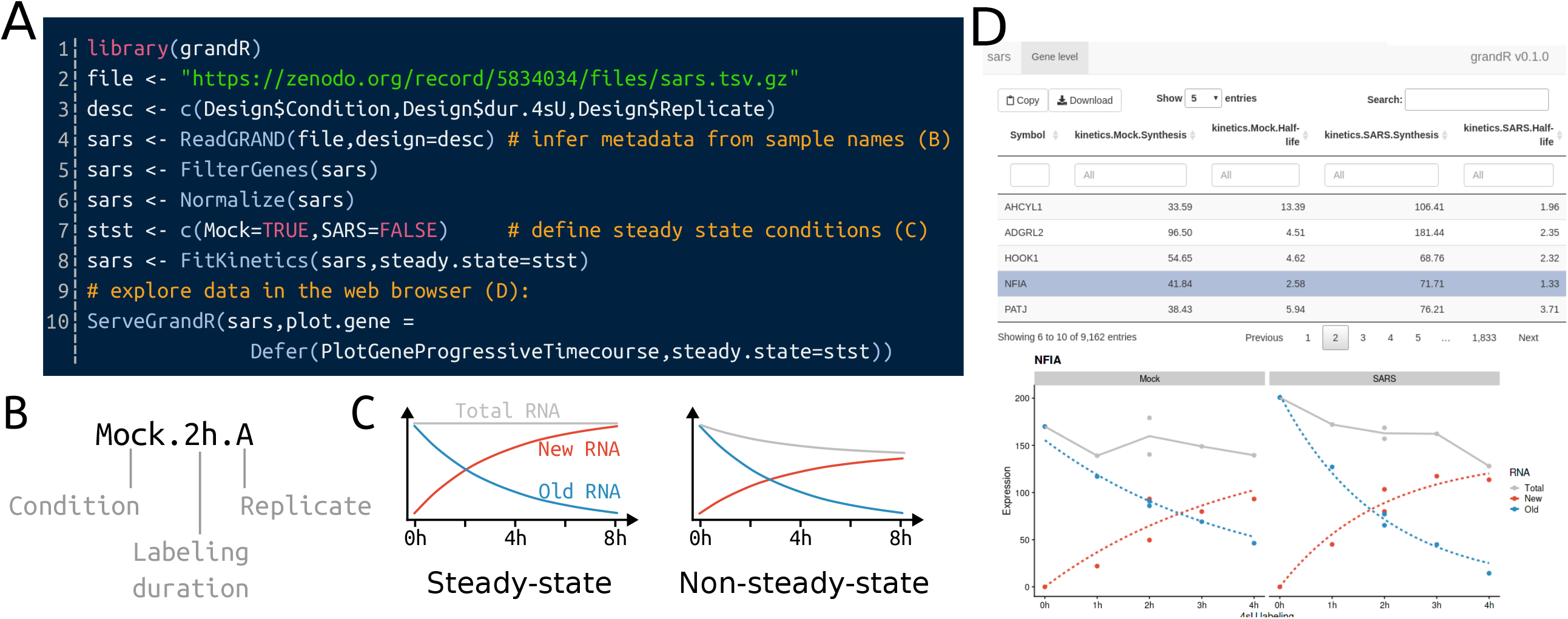
grandR overview **A** Coding example of a grandR project. Self-explanatory high-level commands (blue) load and preprocess data (lines 3-6), and then fit a kinetic model for each gene (line 8). Finally, the interactive web-based tool is started (line 10). Code comments (orange) refer to the other panels. **B** Systematic sample names. When sample names systematically encode metadata in separate fields as shown, grandR can extract these automatically by defining the semantics as shown in lines 3 and 4 in **A. C** grandR can fit kinetic models of RNA with or without assuming steady state expression. **D** Web-based data visualization. This interactive graphical user-interface presents a table of analysis results that can be filtered, searched and exported, and experiment-specific visualizations are displayed for individual genes.

### Quality control reveals impact of long-term labeling >4 h on transcription

For nucleotide conversion approaches, sufficient 4sU incorporation into newly synthesized RNA must be achieved to enable accurate quantification. However, labeling with high 4sU concentrations or labeling over extended periods of time affects cell viability [6] or RNA metabolism [19] in a cell type specific manner. Thus, as opposed to standard RNA-seq, nucleotide conversion sequencing requires additional quality control steps in their analysis workflow.

In grandR, testing for toxicity of 4sU can be performed by comparing 4sU treated samples against equivalent 4sU naïve control samples. To estimate RNA half-lives, Herzog et al. [6] pre-treated mouse embryonic stem cells with low concentrations of 4sU (100μM) for a 24h pulse phase, followed by washing out 4sU and sequencing at several time points during this chase phase. Notably, cell viability was assessed to be ∼80% after 24h. Quality control using grandR revealed that 4sU treated samples and untreated controls segregate in a principal component analysis (Fig. 2A) and that 1,340 out of 8,286 genes were significantly (FDR<5%, DESeq2 Wald test [20]; absolute log_2_ fold change > 0.5 [21]) dysregulated in the 4sU treated sample (Fig. S1A). The p53 pathway was significantly up-regulated, and several stress-related pathways were significantly down-regulated (FDR <0.05, gene set enrichment analysis, Fig. 2B), indicating that central cellular pathways were affected by long-term 4sU treatment. Moreover, RNA half-lives were significantly correlated with the expression changes between 4sU treated and untreated samples (Spearman’s ρ=-0.3, p<2.2×10^−16^, approximate t test; Fig. S1B), indicating that long-term treatment with 4sU affected the RNA metabolism in general. In conclusion, these analyses advocate against long-term 4sU treatment beyond 4h even with low doses of 4sU and, therefore, argue against pulse-chase designs for estimating RNA half-lives in general.

**Figure 2:**
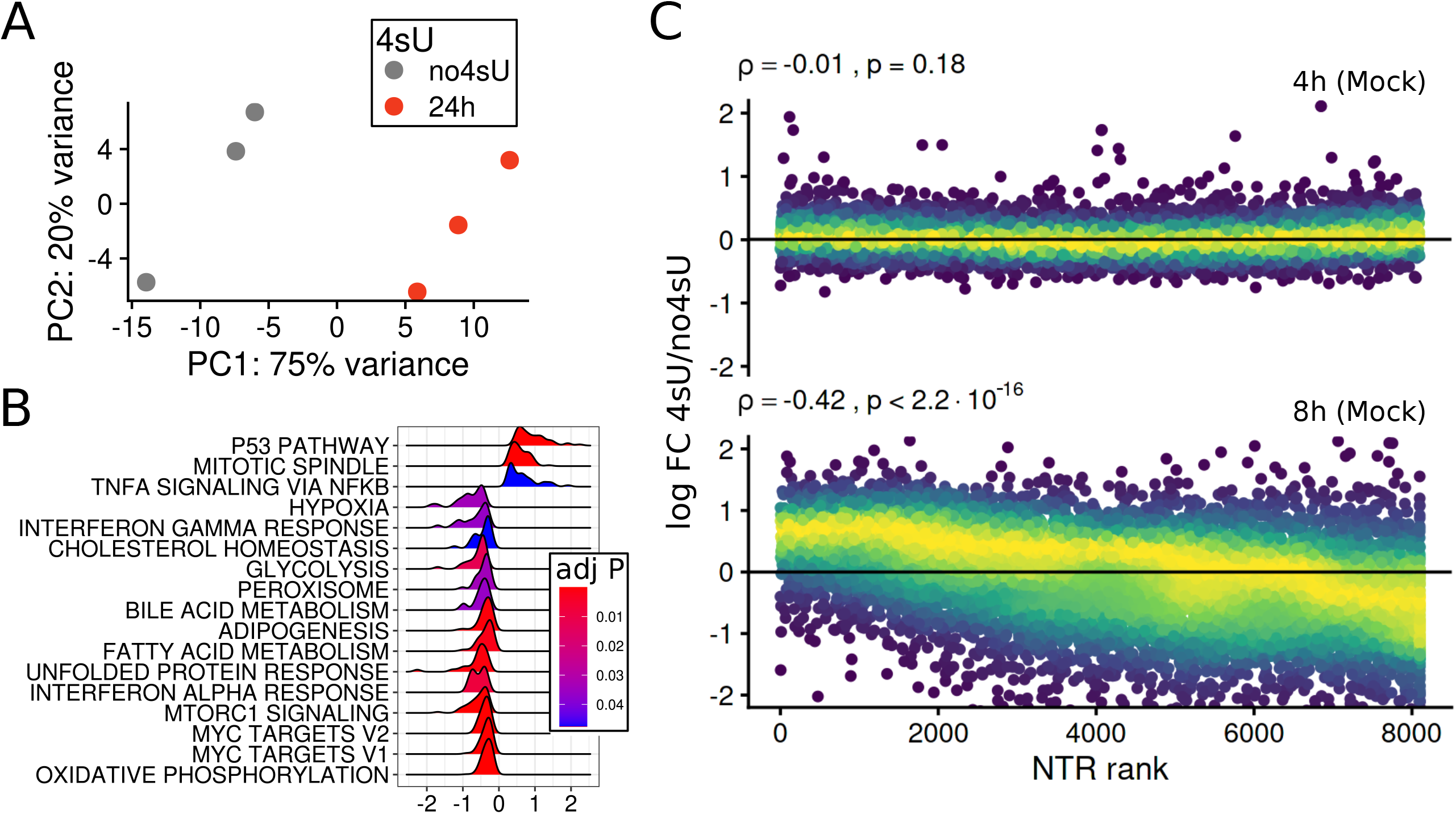
Testing for 4sU toxicity using grandR. **A** Principal component analysis of 3 mouse embryonic stem cell samples treated with 100μM 4sU for 24h and 3 samples without 4sU treatment (no4sU). The percentages of the explained total variance for both principal components shown are indicated. **B** Gene set enrichment analysis of MSigDB hallmark pathways. All pathways with adjusted P values < 5% (Benjamini-Hochberg adjusted for multiple testing) are shown. **C** Scatter plots comparing the ranks of the new-to-total RNA ratios (NTR) of each gene against the log2 fold change of the 4h (upper plot) or 8h (lower plot) sample vs a 4sU naïve sample (untreated mock samples from Ref. [18]). The Spearman correlation coefficients with associated P values (approximate t test) are indicated.

The kinetics of RNA degradation can also be analyzed without chase by monitoring the drop of unlabeled RNA over time. Such a “progressive labeling” design has been used by several studies [1,3,17,18] and provides accurate estimates when timepoints are chosen roughly in the range of the actual RNA half-lives [15,22]. Zuckerman et al [18] performed siRNA knock-down of the nuclear export factor NXF1 and used progressive 4sU labeling for 0h, 2h, 4h and 8h to show that RNA half-lives were not altered. Quality control by grandR revealed that after 8h, but not before, the NTR was significantly correlated with the log_2_ fold change with respect to the (4sU naïve) 0h time point (Spearman’s ρ=-0.42, p<2.2×10^−16^, Fig. 2C). This downregulation of short-lived RNAs in the total RNA pool upon 4sU treatment suggests a general defect in transcription. Importantly for this study the estimated half-lives were not systematically different after excluding the 8h time point from analysis (Fig. S1C). In summary, these analyses indicate that 4sU can substantially impact on transcription before affecting cell viability, and, thus, these effects should be assessed post-hoc in the sequencing data by comparing 4sU treated samples with equivalent 4sU naïve controls.

### Kinetic modeling

The commonly used kinetic model of RNA expression goes back to 1952 [23] and assumes zero-order kinetics for RNA synthesis and first-order kinetics for degradation:

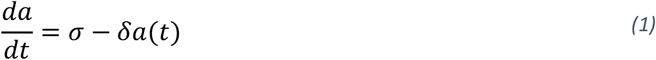

Here, *a*(*t*) is the abundance of RNA at time *t*, and *σ* and *δ* are the rate constants for synthesis and degradation, respectively. A gene is expressed at steady state if 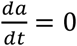, i.e. if 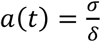. The differential equation (1) can be solved for *a*(0) = *a*_0_:

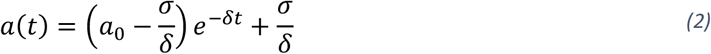

Different variants of this model have been used to estimate RNA stability represented by the degradation rate *δ* or, equivalently, the RNA half-life *t*_1/2_ = log(2) /*δ*: (i) Finkel et al. [17] focused on unlabeled RNA and performed simple linear regression on equation 2 after log transformation and setting *σ* = 0. (ii) In Narain et al. [3], we used non-linear least squares regression (NLLS) to fit the full model in equation 2, which has been done in a similar manner in Zuckermann et al. [18]. (iii) Boileau et al. [24] proposed to fit the full model using their pulseR package [16] based on the raw counts of reads showing T>C mismatches, and to remove bias of this approach using an additional nuisance parameter. (iv) Finally, we have presented a Bayesian method to estimate the degradation rate *δ* under steady state [15].

The main difference among methods (i)-(iv) is the error model employed. While (i) and (ii) assume homoscedastic gaussian errors of estimated pre-existing and newly synthesized RNA levels, either in log space (i) or of the levels directly (ii), pulseR models read counts using a Negative Binomial distribution assuming a gene specific overdispersion parameter that is jointly estimated from all samples. The Bayesian approach (iv) assumes that all data points are generated from a single degradation rate constant *δ* with the only source of error being the random sampling of T>C conversions.

To compare methods (i)-(iv), we implemented *in-silico* simulation of nucleotide conversion sequencing experiments in grandR. Following a previous method to simulate RNA-seq data [25], our simulation procedure first samples read counts from a Negative Binomial distribution with a gene specific overdispersion parameter and expected value both estimated from a real data set. Our simulation then samples T>C mismatches for each individual read based on a user defined NTR value and on sequencing error rates and 4sU incorporation rates estimated from a real data set. Then the simulation estimates NTRs and their posteriors for each gene using the GRAND-SLAM model [15].

We used our read simulator to generate progressive labeling time courses using parameters estimated from a recent SLAM-seq data set of SARS-CoV-2 infection [17] as reference. While the estimated RNA half-lives correlated well with the ground-truth for all methods (R>0.84, p<2.2×10^−16^ for all methods, Pearson correlation; Fig. S2), the non-linear least squared method (ii) and the Bayesian method (iv) were significantly more accurate than the linear regression (i) and pulseR (iii) approaches for data simulated under steady state conditions (Fig 3A). By contrast, non-steady state conditions resulted in generally more substantial deviations from the ground truth (Fig 3B). This analysis also shows that the accuracy of the Bayesian approach (iv) suffers most significantly without steady state. The regression and Bayesian approaches report interval estimates. The regression methods had relatively large confidence intervals, indicating that the gaussian noise approximation does not properly model our simulated sequencing data, especially in logarithmic space as done by method (i) (Fig 3C). The Bayesian method can, very quickly, compute credible intervals by approximating the posterior by a χ^2^ distribution but can also numerically integrate the posterior. For steady state conditions these exact credible intervals were indeed smaller than the approximate intervals (inter-quartile ranges: approximate, [0.37-1.23]; exact, [0.30-0.99]; Fig 3C). Under non-steady state conditions, all deviations were underestimated, most notably for the Bayesian credible intervals, where 88% of the simulated half-lives were outside of the credible interval (Fig S3A). In summary, the non-linear least squares regression (ii) and the Bayesian approach (iv) provided the most accurate estimates under steady state conditions. The Bayesian approach slightly outperformed NLLS, but inherently assumes steady state. We therefore recommend the non-linear least squares regression as the default method for estimating RNA kinetics using progressive metabolic labeling data.

**Figure 3:**
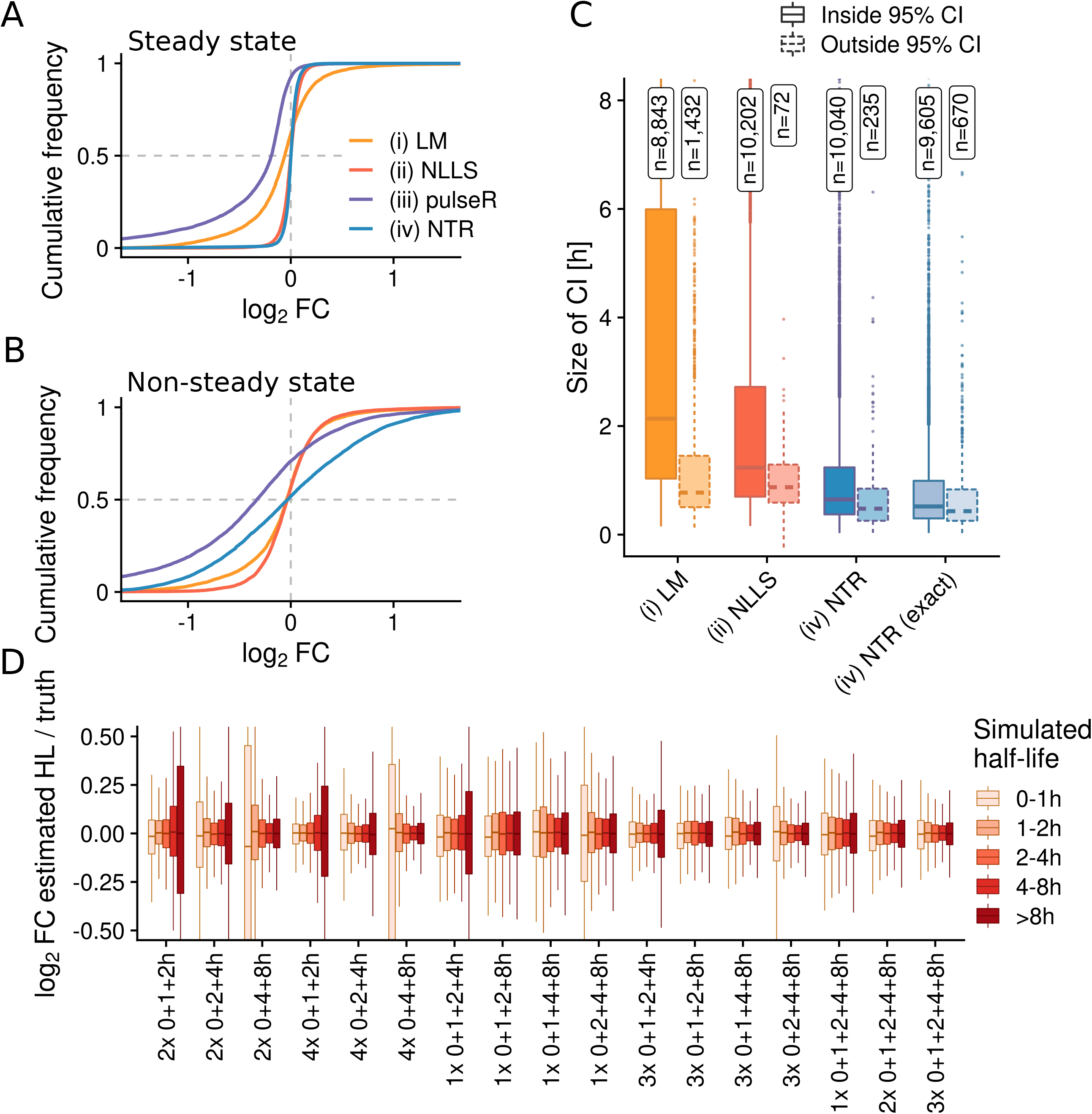
Estimating half-lives using progressive labeling experiments. **A-B** Empirical cumulative distributions of log2 fold changes of estimated half-lives vs. ground-truth for the linear model (LM), the non-linear least squares method (NLLS), the pulseR method and the Bayesian approach (NTR). In **A**, the ground-truth is simulated under steady state conditions, in **B** the simulation starts from an initial value *a*_0_ ≠ *σ*/*δ* for each gene (see Methods). **C** Boxplots showing the sizes of 95% half-life confidence intervals (CI; for LM and NLLS) or 95% half-life credible intervals (CI; for NTR). Simulations were performed under steady state conditions. Distributions for genes having the ground-truth inside or outside of the estimated CI are shown separately and the numbers of these genes are indicated. NTR represents the χ^2^ approximation of CIs, NTR (exact) represents exact CIs computed numerically. **D** Boxplots showing log2 fold changes of half-lives estimated by the NLLS method vs the ground truth of simulated data under steady state conditions. The distributions for different half-life classes are shown for several experimental settings involving the indicated number of replicates and time points.

### Choosing number of replicates, time points and sequencing depth

Our simulation also enabled us to assess how many reads, replicates and time points are required to obtain accurate estimates of kinetic parameters. We reasoned that a moderate number between 6 and 12 samples per condition should be used. However, it is a priori unclear whether these should be distributed over many time points, or whether more replication of the same time points is more beneficial. We therefore simulated a broad range of potential experimental settings (Fig 3D). As expected from our previous analyses [15], if early (1h) or late (8h) time points are missing, the estimates for short-lived or long-lived RNAs, respectively, suffer significantly. This became most obvious when we only simulated a single time point (Fig S3B). Thus, to analyze the complete landscape of RNA half-lives multiple time-points spanning the whole range of RNA half-lives are required. We next simulated data for a full progressive time course (1h, 2h, 4h and 8h) with different sequencing depths and numbers of replicates (Fig S3C). Interestingly, increasing the number of replicates per time point boosted the accuracy stronger than increasing the number of reads. In conclusion, our data show that time-points must be carefully chosen and the costs for sample and library preparation must be weighed against the sequencing costs to obtain accurate RNA half-lives.

### Temporal recalibration improves the model fit

4sU is not available for transcription immediately once the cells are cultured on 4sU media, but it is actively transported across cell membranes via nucleoside transporters and is processed by the pyrimidine salvage pathway before it is available as substrate for transcription. Thus, the concentration of active 4sU increases until saturation, and RNA that was transcribed significantly before reaching saturation contains fewer 4sU than RNA transcribed later. Therefore, especially for earlier timepoints, the effective labeling time is expected to be much shorter than the nominal labeling time.

To test this effect, we used grandR to estimate RNA half-lives for published data of Calu-3 cells infected with SARS-CoV-2 and mock infected control cells [17]. Indeed, the residuals of the model fit were mostly negative for the 1h time points, and more balanced at later time points (Fig. 4A-B). Since the effective labeling time for a sample is a global parameter that is common to all genes, and new and old RNA follow the model defined in equation (2) for all genes, we reasoned that it should be possible to estimate effective labeling times by maximizing the joint likelihood of all gene specific synthesis and degradation rates and the effective labeling times.

**Figure 4:**
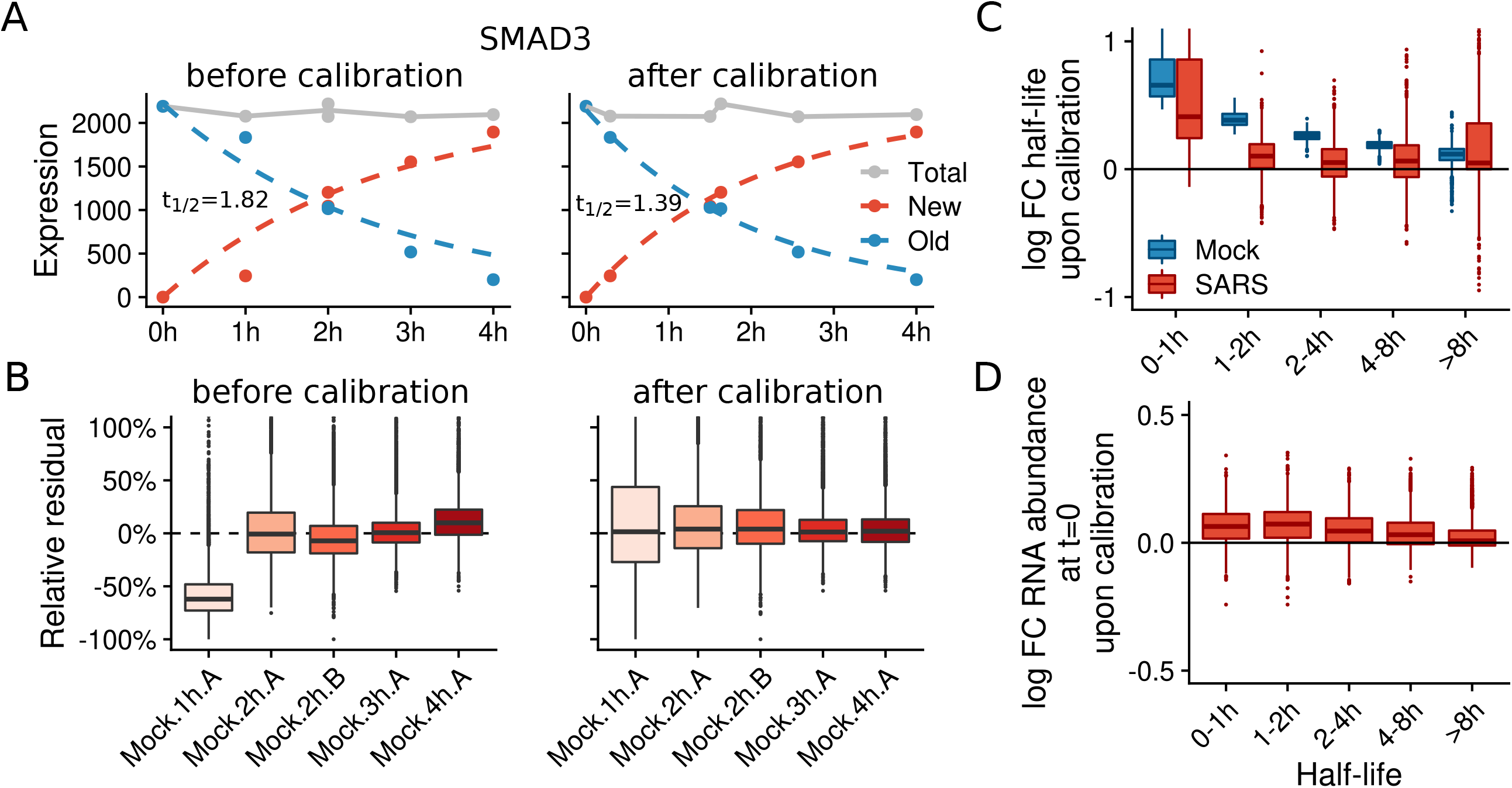
Temporal recalibration of SARS-CoV-2 SLAM-seq data. **A** Progressive labeling plots of the SMAD3 gene before (left) and after (right) temporal recalibration. Points represent the total, new or old read count of SMAD3 at the indicated time after labeling. Dashed lines show the model fit (NLLS method). Estimated half-lives are indicated. **B** Boxplots showing relative residuals from the model fit (NLLS method) before (left) and after (right) temporal recalibration of n=9,162 genes for all samples. **C-D** Boxplots showing log2 fold changes (recalibrated vs. uncalibrated) of half-lives for the mock and virus infected (SARS) samples (**C**) or of the estimated initial abundances *a*_0_ (recalibrated vs. uncalibrated) for the non-steady state infected samples (**D**) for n=9,163 genes. Separate distributions for genes from different half-life classes are shown.

We first tested this temporal recalibration by simulated time courses where we artificially changed the nominal labeling times. The recalibrated labeling times were on average within 1.03-fold of the true effective labeling time (Fig. S4A), and estimation of kinetic parameters was completely rescued after recalibration (Fig. S4B). We then recalibrated the labeling times for the SARS-CoV-2 data. Indeed, the residuals became smaller for all samples and were now symmetric (Fig. 4A-B). Globally, temporal recalibration affected short half-lives stronger than long half-lives (Fig. 4C), presumably because early time points are the most informative to estimate short half-lives. Moreover, for the virus infected samples there were substantially more gene-specific differences at the 1h time point, indicating that without steady state assumption, the first time point is important to estimate the initial abundance *a*_0_. Indeed, the estimates of *a*_0_ exhibited the same gene specific differences like half-life estimates upon calibration (Fig. 4D). In conclusion, due to the kinetics of 4sU uptake and activation, the effective labeling time might differ from the nominal labeling time, especially for short labeling. For progressive labeling experiments, this can be corrected by temporal recalibration.

### Estimating changes in synthesis or degradation from snapshot experiments

Nucleotide conversion sequencing has also applications beyond progressive labeling time courses. We and others showed that new RNA from a single “snapshot” timepoint is more sensitive to detect short-term changes of gene expression than standard RNA-seq without metabolic RNA labeling, e.g. upon virus infection [11], drug treatment, or acute depletion of transcription factors via degron systems [3,26]. So far, analyses of snapshot samples have been performed in an ad-hoc manner by the application of standard differential gene expression tools on estimated new or old RNA.

We have previously shown that steady-state half-lives can, in principle, be estimated from a single snapshot sample [15]. Here, we extend this and show that both synthesis and degradation rates (*σ* and *δ*) can be estimated also without assuming steady state:

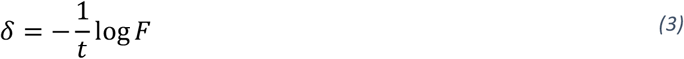

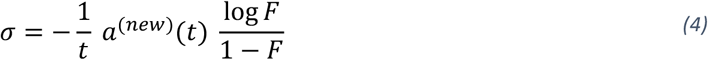

Here, *a*^(*new*)^(*t*) is the abundance of new RNA at time *t*, and 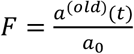is the ratio between pre-existing (old) RNA *a*^(*old*)^(*t*) at time *t* and the total level *a*_0_ at time 0. Thus, to compute *σ* and *δ*, in addition to old and new RNA, *a*_0_ must be known, either due to the assumption of steady state (where 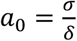), or by a separate sample measured at time *t* = 0. Importantly, equations 3 and 4 can be extended to enable computing *σ* and *δ* when a separate sample is available for any time *t* < 0 (see Methods).

Estimates of *σ* and *δ* based on applying equations 3 and 4 to measured data might be highly inaccurate due to the NTR quantification uncertainty, due to a labeling time not matching the gene specific RNA half-life, or sampling noise due to low numbers of reads. In addition to these technical factors, *σ* and *δ* are also subject to biological variability among replicate samples. To control these factors, we developed a Bayesian hierarchical model to estimate the joint posterior distribution of *σ* and *δ* as well as the joint posterior of 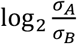 and 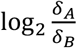 for differential analysis of two samples A and B.

To test our approach, we simulated data for two conditions with 2h labeling. We left one condition at steady state, for the other we either perturbed synthesis or degradation rates, or left them unperturbed as control. The maximum-a-posteriori log fold change estimates of both *σ* and *δ* were unbiased, and much more accurate for *σ* (root mean square deviation (RMSD) = 0.048; Fig. 5A) than for *δ*(RMSD = 0.510; Fig. 5B). Estimated changes in *σ* reflected the true change in synthesis rates more accurately than new RNA (RMSD = 0.064; Fig. 5C). Counterintuitively, the old RNA fold change did not correspond to the true fold change of RNA half-lives (Fig. 5D). Indeed, equation 3 shows that an observed fold change of old *RNA f between* two conditions A and B means that the degradation rates differ by the additive constant − 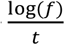 rather than a multiplicative factor. Previously, an observed new RNA fold change has been equated with a change in synthesis rate [14]. However, equation 4 shows that new RNA fold changes are also affected by changes in degradation rates, predominantly for genes with short-lived RNAs. Indeed, we observed significant changes in new RNA when only the degradation but not the synthesis rates were changed, which was restricted to genes with short-lived RNAs (Fig 5E). Of note the estimated synthesis rate changes by our Bayesian model were not affected by changes of RNA stability. For unperturbed controls, estimated changes of *δ* exhibited more variance than estimated changes of *σ*. This effect was much less pronounced for genes with short RNA half-lives, or when the labeling duration was 4h instead of 2h (Fig. 5F).

**Figure 5:**
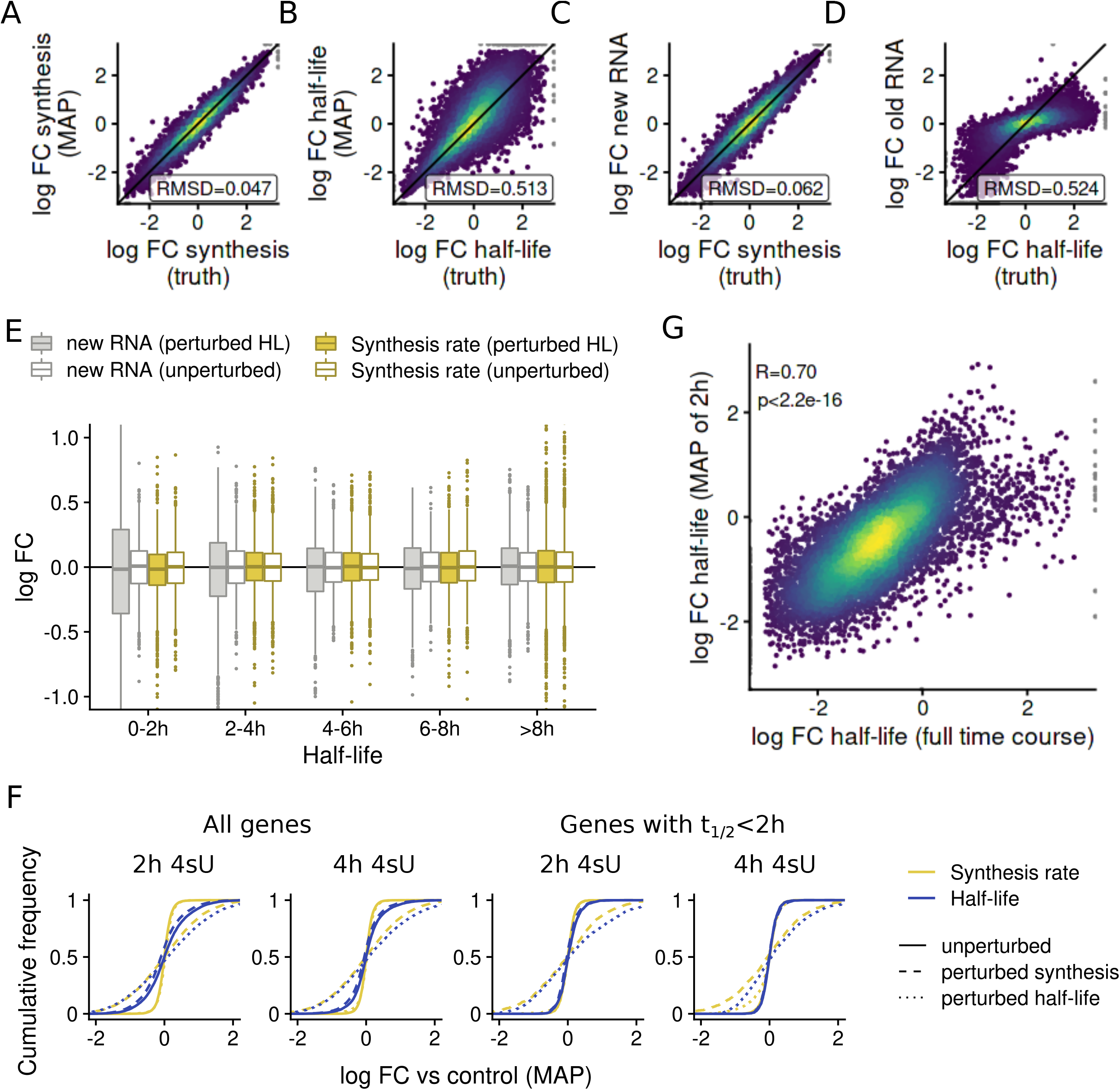
Estimating changes in synthesis or degradation from snapshot experiments. **A-D** Scatterplots comparing simulated log2 fold changes against maximum-a-posteriori (MAP) estimates of RNA synthesis log2 fold changes (**A**), MAP estimates of RNA half-life log2 fold changes (**B**), observed new RNA log2 fold changes (**C**) or old RNA log2 fold changes (**D**). Three replicates at 2h of labeling were simulated after perturbing synthesis (**A**,**C**) or half-lives (**B**,**D**) for 2h and compared against unperturbed controls. The root mean square deviations (RMSD) over all n=10,835 simulated genes are indicated for each comparison. **E** Boxplots showing the log2 fold changes of new RNA or of estimated synthesis rates either for the simulated samples with perturbed RNA half-lives (perturbed HL) or the unperturbed samples vs the controls. Separate distributions for genes from different simulated half-live classes are shown as indicated. **F** Empirical cumulative distributions showing log2 fold changes of either estimated synthesis rates (yellow) or RNA half-lives (blue). For each distribution, either unperturbed samples (solid lines), samples with perturbed synthesis rates (dashed lines) or samples with perturbed half-lives (dotted lines) were compared against controls. Distributions are shown for all genes, only for genes with short RNA half-lives t_1/2_<2h, and for simulated labeling of 2h or 4h, as indicated. **G** Scatterplot comparing log2 fold changes of RNA half-lives estimated from the full progressive labeling time courses using the NLLS method (x axis) or the MAP estimator from our Bayesian model using the 2h time point only. The Pearson correlation and the associated P value (approximate t test) are indicated.

We then applied our Bayesian model for changes of RNA stability to the 2h time point of the SARS-CoV-2 data [17], revealing that the degradation rate changes recapitulated the changes identified by modeling the full progressive labeling time course (R=0.7, p<2.2×10^−16^, Pearson correlation; Fig. 5G).

In summary, in contrast to previously used fold changes of old and new RNA, the maximum-a-posteriori estimates of our hierarchical model provide directly interpretable log fold changes of synthesis and RNA half-lives from snapshot data.

### ROPE analysis of significant changes of *σ* and *δ*

We analyzed “regions of practical equivalence” (ROPE) [27] to quantify significant changes of synthesis or degradation using our Bayesian approach. As a measure of significance, we used the posterior probabilities *P*_*σ*_ or *P*_*δ*_ of the log_2_ fold change (synthesis or degradation, respectively) being either less than − 0.25 or greater than 0.25. As a comparison, we analyzed Benjamini-Hochberg adjusted P values *q*_*new*_ and *q*_*old*_ computed by DESeq2 [20] for new and old RNA, respectively.

We first analyzed our simulated data (2h labeling) where RNA synthesis rates were perturbed. As expected from overall n=10,835 genes, virtually none had *P*_*δ*_ > 0.9. (n=127, 1.2%) or *q*_*old*_ < 0.01 (n=0) independent of the true change of RNA synthesis (Fig 6A). Notably, of the n=3,388 genes simulated to be more than 2-fold up- or down-regulated, n=3,249 (95.9%) and n=3,174 (93.7%) genes had *P*_*σ*_ > 0.9 and *q*_*n*_ < 0.01, respectively (Fig. 6A). Thus, ROPE analysis of our Bayesian model and DESeq2 analysis of new and old RNA showed similar sensitivity and specificity when only RNA synthesis rates are changed.

**Figure 6:**
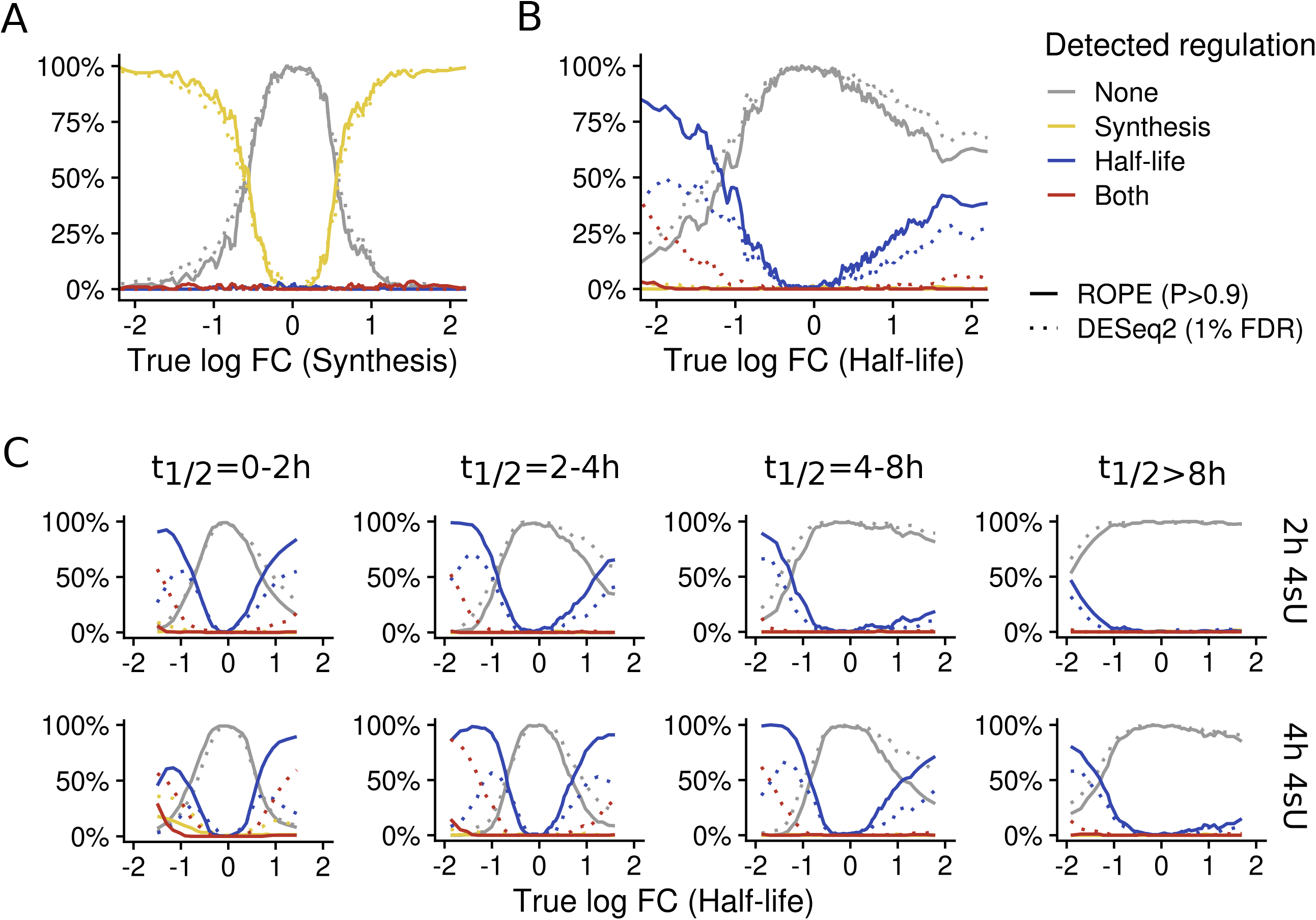
Region of practical equivalence analysis of simulated data. Line plots comparing two criteria for differential regulation of synthesis rates (A) or half-lives (B-C) are shown. For **A**, the two criteria are the ROPE probability for synthesis *P*_*σ*_ > 0.9 and the DESeq2 P value (Wald test, Benjamini-Hochberg multiple testing adjusted) for new RNA *q*_*n*_ < 0.01. For **B** and **C**, the two criteria are the ROPE probability for degradation *P*_*δ*_ > 0.9 and the DESeq2 P value (Wald test, Benjamini-Hochberg multiple testing adjusted) for old RNA *q*_*o*_< 0.01. The x axis represents rolling statistics (bin width 200 genes) over the log2 fold change of synthesis rates (**A**) or RNA half-lives (**B**) for the simulations with perturbed synthesis and half-lives vs. control, respectively. The different lines show the percentage of genes in a bin with detected regulation in synthesis, half-life, both or none. **A** and **B** show all genes for 2h of 4sU labeling. **C** shows genes of different half-life classes and for 2h or 4h of 4sU labeling, as indicated.

Next, we focused on simulated data where RNA half-lives were perturbed. From overall n=10,835 genes, n=40 (0.4%) had *P*_*σ*_ > 0.9 and n=407 (4.0%) had *q*_*new*_ < 0.01 (Fig 6B). These hundreds of genes with significant changes in new RNA predominantly had downregulated RNA half-lives (n=302 out of overall 1641 genes with >2-fold downregulated RNA half-lives, 18.4%). Thus, as shown above, new RNA for some genes exhibited significant changes when only RNA half-lives are changed. By contrast, our Bayesian approach can accurately differentiate between changes in synthesis and degradation. Moreover, out of n=3,390 genes with >2-fold regulated half-lives, n=1,692 (49.9%) had *P*_*δ*_ > 0.9 and n=1,351 (39.8%) had *q*_*old*_ < 0.01 (Fig 6B), indicating that our Bayesian approach is also more sensitive than analyzing old RNA for detecting changes in RNA stability.

Interestingly, the sensitivities of our Bayesian approach for detecting changes in RNA stability were asymmetric (65.7% for downregulated RNA half-lives and 33.0% for upregulated RNA half-lives; Fig 6B). To investigate this further, we stratified genes according to their unperturbed RNA half-lives. For genes with RNA half-lives of less than 2h, sensitivities indeed were symmetric, but exhibited increasing asymmetry for longer half-lives (Fig 6C). This asymmetry can be explained by the fact that half-lives not matching to the duration of 4sU labeling cannot be estimated accurately. To corroborate this, we repeated these analyses with 4h of simulated 4sU labeling, which resulted in symmetric sensitivities for average half-lives of 2-4h. In summary, our Bayesian approach can accurately differentiate between effects on RNA synthesis and degradation.

### Bayesian analysis indicates target gene specific differences of regulation by acute BANP depletion

We utilized our Bayesian modeling approach for the analysis of published data from cells after degron-mediated depletion of BANP, which has recently been revealed to bind to unmethylated CGCG motifs in CpG islands to promote transcription of a set of essential genes [26]. For this study, samples from multiple timepoints (1h, 2h, 4h, 6h and 20h) after depletion of BANP were labeled with 4sU prior to sequencing. Importantly, the samples from the 4h timepoint and later were labeled for 2h, but shorter labeling of 30 and 90 minutes was applied for the 1h and 2h timepoints, respectively. Due to these different labeling times, new RNA is not directly comparable among the samples and inference of *σ* is required to interpret the data. We first calibrated labeling times, which can here be accomplished by matching the transcriptome-wide distribution of degradation rates. Indeed, after recalibration, the distribution for RNA half-lives estimated for each timepoint were largely indistinguishable (Fig S5A), and the same was also true for RNA synthesis rates (Fig S5B).

For each timepoint, the RNA synthesis log fold changes for BANP targets determined by ChIP-seq [26] were significantly and consistently shifted towards negative values compared to non-targets (Fig. 7A). This is remarkable especially for the 1h timepoint, where 4sU labeling only was 30 minutes, and suggests that synthesis rates were reduced immediately once BANP was depleted from cells, and then stayed constant for at least 20h. To further investigate this, we analyzed the synthesis log fold change posterior distributions for individual genes. This revealed that there were substantial gene specific differences, with some BANP targets like Taf1d showing gradually decreasing synthesis rates with efficient downregulation only later than the 1h timepoint (Fig 7B) and for others like Herc1 (Fig 7C) or Tupgcp5 (Fig 7D) synthesis rates dropped early and rose again later suggesting negative feedback loops. In conclusion, the Bayesian hierarchical model implemented in grandR can be used to uncover detailed information about gene regulation from snapshot experiments reflecting genome-wide trends and to generate testable hypotheses of individual genes.

**Figure 7:**
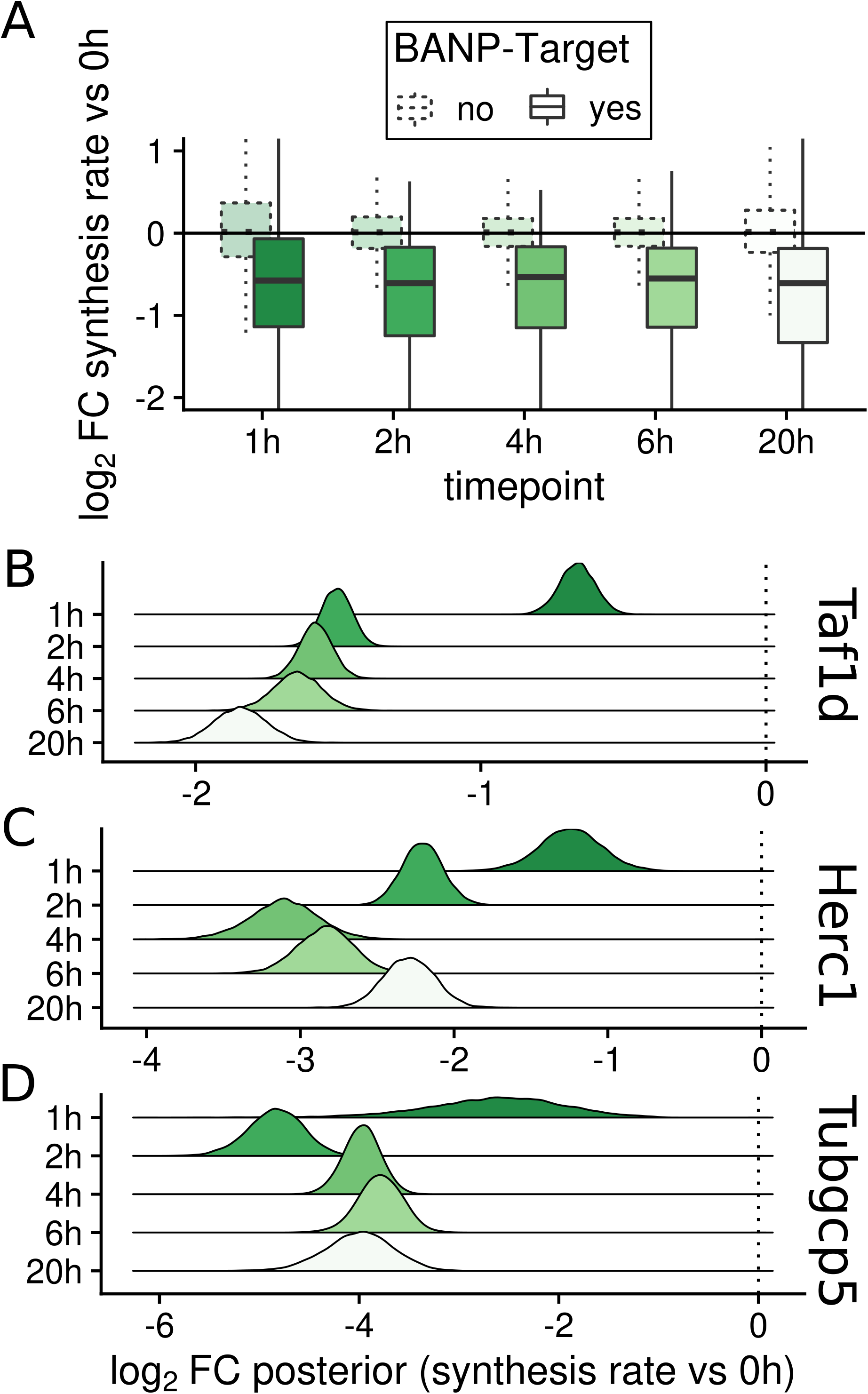
Dynamic regulation of synthesis rates upon acute BANP depletion. **A** Boxplots showing log2 fold changes of synthesis rates for several experimental time points vs the 0h time point. Distributions of BANP target genes (according to ChIP-seq experiments) are shown separately from non-target genes. **B-D** Estimated posterior densities for Taf1d (**B**), Herc1 (**C**) and Tubgcp5 (**D**) of log2 fold changes of the synthesis rates of the indicated time points vs. the 0h time point.

## Discussion

Nucleotide conversion sequencing is now widely used to infer kinetic parameters of RNA expression but there is still a lack of computational tools for data analysis. Our goal for developing grandR was to provide a comprehensive and easy-to-use toolkit to facilitate a broad range of different analysis steps for such data.

It is well described in the literature that nucleoside concentrations must be optimized for specific cell types and desired labeling times [5,6,19]. Toxicity of too highly concentrated 4sU has previously been assessed by testing for cell viability. However, here we show that transcription can be affected before cell viability suffers. We therefore advocate that all studies employing metabolic RNA labeling must report the extent of any effect of 4sU on transcription. Since levels of short-lived RNAs quickly decline when transcription is globally inhibited, a correlation of the fold changes for 4sU treated samples vs. corresponding 4sU naïve controls with the NTRs can be used as surrogate marker for transcriptional defects due to 4sU treatment, as implemented in grandR.

RNA degradation rates have previously been estimated using progressive labeling time courses using different computational methods. All these approaches employed the kinetic model described by equations 1 and 2 but they differ in their choice of the error model. Due to noise introduced by the inference of the NTR, the actual errors of normalized new and old RNA likely are differently distributed than standard RNA-seq data. Our simulations indicate that the errors are well approximated by a gaussian distribution. We recommend using the non-linear least squares fitting procedure as the general tool for fitting the kinetics of RNA expression. The Bayesian approach provides slightly more accurate results and better error bounds but can only be used under steady state conditions.

We have also shown that much simpler snapshot experiments can be used to infer the kinetics of RNA expression. A major advantage of such snapshots is that, instead of using multiple time points to observe and characterize the drop of pre-existing RNA for obtaining the degradation kinetics, with the same number of samples the kinetics can be probed at multiple time points, e.g. after virus infection. This is of particular importance when synthesis or degradation rates are not constant. Indeed, the reduced RNA half-lives observed for SARS-CoV-2 likely result from the general host shutoff protein nsp1 encoded by SARS-CoV-2 [28]. It is therefore very likely that degradation of cellular mRNAs depends on the abundance of nsp1, which increases substantially over the first few hours of infection. Thus, the degradation rate at 4h post infection (corresponding to the 1h labeling time point in the SLAM-seq data from Ref. [17]) likely is different at 7h post infection (the final 4h labeling time point).

A major caveat of metabolic RNA labeling experiments is that the effective labeling time might not correspond to the nominal labeling time. grandR provides tools to test for this critical issue and recalibrate labeling times: For progressive labeling time courses, asymmetric residuals of early time points indicate shorter effective labeling times. Using the labeling times as additional independent variables when jointly fitting the kinetics for all genes, as implemented in grandR, can be used to estimate the effective labeling times. For snapshot experiments, labeling times can be recalibrated based on additional assumptions, e.g. based on reference RNA half-lives. Testing for effective labeling is critical when samples with distinct labeling times are compared, and for estimating RNA degradation and, to a lesser extent, synthesis rates in absolute terms.

## Conclusions

Nucleotide conversion approaches greatly reduced the burden on the wet-lab side for conducting metabolic RNA labeling experiments but introduced the need for more sophisticated tools for their computational analysis. Complementing our GRAND-SLAM software for primary processing of such data, we developed the grandR package as a general toolkit to aid researchers to further analyze and interpret such data. Here, we demonstrated that additional quality control measures are necessary for such data to exclude effects of 4sU on transcription, and that short labeling times often require recalibration. For both methods, grandR provides high-level functions. Furthermore, grandR enables researchers to estimate synthesis and degradation rates for both progressive labeling as well as snapshot experiments without requiring steady state assumptions. Finally, grandR provides a web-based interface for exploratory data analysis.

## Methods

### SLAM-seq preprocessing

All SLAM-seq data used here were processed using the GRAND-SLAM pipeline [15]. Fastq files were downloaded from the SRA database. The accession number were: GSE99970 for the 24h 4sU labeling data set from Ref. [6] (samples: GSM2666816, GSM2666817, GSM2666818, GSM2731767, GSM2731768, GSM2731769), GSE139151 for the NXF1 knockdown data set from Ref. [18], GSE162323 for the SARS-CoV-2 data set from Ref. [17], and GSE155604 for the BANP depletion data set from Ref. [26]. Adapter sequences were trimmed using cutadapt (version 3.4) for the SARS-CoV-2 data, where reads were not pre-trimmed on SRA. Then, bowtie2 (version 2.3.0) was used to map read against an rRNA (NR_046233.2 for 24h and BANP, and U13369.1 for NXF1 and SARS-CoV-2) and Mycoplasma database. Remaining reads were mapped against target databases using STAR (version 2.5.3a). We used the murine genome for 24h and BANP, the human genome for NXF1, and the combined human and SARS-CoV-2 (NC_045512) genome for SARS-CoV-2. All genome sequences were taken from the Ensembl database (version 90). Bam files for each data set were merged and converted into a CIT file using the GEDI toolkit [29] and then processed using GRAND-SLAM (version 2.0.7; for the NXF1 data set, GRAND-SLAM 2.0.5g was used).

### Read simulation

To simulate a nucleotide conversion sequencing experiment for *n* genes with relative abundances *a*_1_, …, *a*_*n*_, ∑_*i*_ *a*_*i*_ = 1, overdispersion parameters *d*_*i*_ and total read count *N*, first *n* random numbers *C*_*i*_ are drawn from negative binomial distributions *NegBinom*(*a*_*i*_ *N, d*_*i*_). We use the parametrization *NegBinom* (*μ, d*) such that the mean is *μ* and the variance is *μ*+ *dμ*^2^.

To simulate the “measured” NTR for gene *ntr*_*i*_ given the true NTR *t*, we sampled the number of uridines *u*_*r*_ covered by each of the *c*_*i*_ reads from a binomial distribution *Binom* (*rl, p*_*u*_), where *rl* is the user-defined read length (used here: 75), and the probability for an uridine at any position *p*_*u*_ is sampled from a beta distribution with user-defined average uridine content (used here: 0.25) and standard deviation thereof (used here: 0.05). For each read *r*, then the number of conversions *tc*_*r*_ is sampled from a Binomial mixture distribution *BinomMix* (*u*_*r*_, *p*_*e*_, *p*_*c*_, *ntr*_*i*_) defined by the probability function

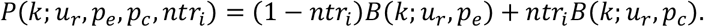

Here, 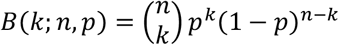 is the probability function of the Binomial distribution. We used a sequencing error rate of *p*_*e*_ = 10^−4^ and a T-to-C conversion rate of *p*_*c*_ = 0.04. Of note, *D*= (*u*_*r*_, *tc*_*r*_), *r* ∈ {1, …, *c*_*i*_} for a gene *i* represent the sufficient statistics for the GRAND-SLAM model. Then, GRAND-SLAM is used to obtain the maximum-a-posteriori (MAP) estimate for the NTR by numerically maximizing the Binomial mixture log likelihood function [15], i.e. we used a uniform prior. In addition, it also computes the Beta approximation of the posterior distribution of *ntr*_*i*_ as described [15].

This procedure is implemented in the function *SimulateReadsForSample* of grandR and simulates the read count *C*_*i*_ and the NTR Ξ_*i*_ for *n* genes based in several user-defined parameters as described above. grandR also provides the higher-level function *SimulateTimeCourse*. Based on a time point *t* RNA synthesis rates *σ*_*i*_, degradation rates *δ*_*i*_, initial abundances *a*_0*i*_ and global synthesis and degradation variance parameters *v*_*σ*_ and *v*_*δ*_ (here both were set to 1.05), this function computes both the relative abundances *a*_*i*_ and the new-to-total RNA ratio *ntr*_*i*_, i.e. the parameters to *SimulateReadsForSample* as follows: To model biological variability, we define 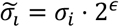 where *ϵ* is gaussian noise. Here, the noise level was chosen such that the 95% quantile of the gaussian is equal to log_2_ *v*_*σ*_. This way, 90% of all 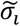 are expected to be at most *v*_*σ*_-fold less or greater than *σ*_*i*_. Equivalently, we defined 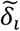. Then, the abundance of new and old RNA at time *t* is computed as 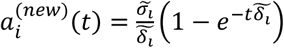 and 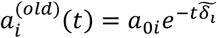 respectively. Thus, 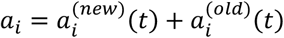 and 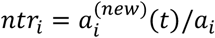.

For all simulations here, we used the “mock” samples from the SARS-CoV-2 data set to compute the relative abundances *a*_*i*_ and estimated the overdispersion parameters *d*_*i*_ using the function *estimateDispersions* from the DESeq2 package. The reference synthesis rates *σ*_*i*_ and degradation rates *δ*_*i*_ were estimated using the NLLS approach from the same data set. To simulate perturbed synthesis rates 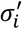, degradation rates 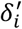 or initial abundances 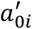 (to start from non-steady state conditions), we sampled gaussian noise such that ∼5% of all genes are expected to be perturbed at most 2-fold.

### Kinetic model

To model the abundance of RNA at time *t, a*(*t*), we use equation 2:

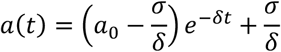

Here, *σ* and *δ* are the rate constants for synthesis and degradation, respectively, and *a*_0_: = *a*(0) is the initial abundance at time *t* = 0. We define the following two functions for the abundances of old and new RNA, respectively, after labeling for time *t*:

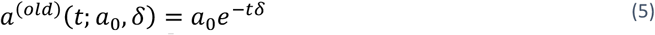

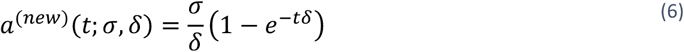

Under steady state assumptions, we have 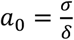 and can use the steady state function instead of equation 5:

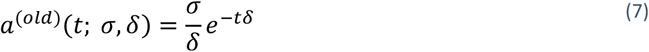

We have data given in the form of total expression values *C*_*k*_ and “measured” NTRsΞ_*k*_ for samples taken at time points *t*_*k*_. Note that even if we use the same notation *C* as for read counts above in “Read simulation”, here we assume that *C*_*k*_ is a normalized expression measure. We further index by *k* to indicate the biological samples where data were obtained and drop the gene index *i* for clarity. The NTRsΞ_*k*_ actually are not measured but are estimates of the parameter 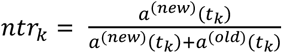. We also have the beta approximation of the posterior distribution of *ntr*_*k*_ given by *α*_*k*_ and *β*_*k*_, i.e. *ntr*_*k*_ | *D*∼ *Beta*(*α*_*k*_, *β*_*k*_) for data *D*. We use bold face ***t*** = (*t*_1_,… *t*_*m*_), ***α***= (*α*_1_, … *α*_*m*_), ***β***= (*β*_1_,… *β*_*m*_), etc. to denote the vectors valued parameters.

### Modeling progressive labeling time courses

We define the random variables for old and new RNA as *O*_*k*_ = *C*_*k*_ ·(1− Ξ_*k*_) and *N*_*k*_ = *C*_*k*_· Ξ_*k*_. The distributions of *N*_*k*_ and *O*_*k*_ dependent on measurement noise from the sequencing experiment, uncertainty in the estimate Ξ_*k*_ and biological variability. In a “progressive” labeling experiment each sample is labeled for a duration of *t*_*k*_ starting from a common time point *t* = 0, and *σ* and *δ* from equations 1 and 2 remain constant after time *t* = 0. Then the expected values of *O*_*k*_ and *N*_*k*_ are

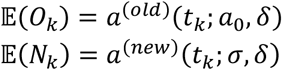

All methods described below, except for pulseR, can conveniently be used in grandR using the function *FitKinetics*, which will fit either the LM, the NLLS or NTR model for each gene by calling the respective functions mentioned below.

### LM method

For the linear model approach (method (i) in the text), we note that we have a linear function log (*a*^(*old*)^(*t*; *a*_0_, *δ*)) = log (*a*_0_)− *δt* after log transforming equation 5. Thus, *δ* and *a*_0_ can be estimated using simple linear regression. Under the assumption of steady state, we can also obtain an estimate of *σ* = *a*_0_· *δ* Note, however, that this assumes all *O*_*k*_ to follow homoscedastic LogNormal distributions. We deem this model quite unrealistic, as at late time points *t*_*l*_ ≫ *t*_1/2_ (where *t*_1/2_ = log(2) /*δ* is the half-life), *a*^(*old*)^(*t*_*l*_ ; *a*_0_, *δ*) quickly approaches 0, and we therefore expect the residual log (*a*^(*old*)^(*t*_*l*_; *a*_0_, *δ*))− log(*O*_*l*_) to be far greater than log (*a*^(*old*)^(*t*_*e*_ ; *a*_0_, *δ*))− log(*O*_*e*_) at an earlier time point *t*_*e*_ < *t*_1/2_. This approach is implemented by the *FitKineticsGeneLogSpaceLinear* function in grandR using the *lm* function of R. Confidence intervals are estimated using the *confint* function.

### NLLS method

For the non-linear least squares approach (method (ii) in the text), we assume *O*_*k*_ and *N*_*k*_ to be homoscedastic gaussian. Thus, *σ, δ* and *a*_0_ (or *σ* and *δ* under steady state assumptions) can be estimated using non-linear least squares regression. This is implemented in grandR by the function *FitKineticsGeneLeastSquares* using the *nls*.*lm* function from the minpack.lm package. Confidence intervals are estimated using *confint*.*nls*.*lm*.

### pulseR method

pulseR (method (iii) in the text) originally was developed for 4sU labeling experiments where labeled and unlabeled RNA was physically purified and sequenced separately [16]. It was later adapted to also handle nucleotide conversion sequencing data [24]. pulseR operates on labeled and unlabeled read counts (i.e. reads with and without observed T-to-C conversions), and includes additional nuisance parameters to model reads from unlabeled RNA with T-to-C conversions (e.g., sequencing errors) and reads from labeled RNA without T-to-C conversions (reads not covering 4sU incorporation sites). In our notation, the pulseR model is

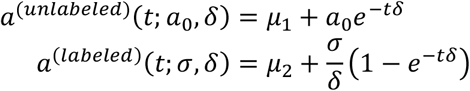

Here, *μ*_1_ is the fraction of reads without T-to-C conversion, that indeed is not derived from old RNA, and *μ*_2_ is the fraction of reads with T-to-C conversions, that indeed is not derived from new RNA. Parameters are estimated using the counts of reads with and without T-to-C conversions instead of estimated old and new RNA levels *O*_*k*_ and *N*_*k*_ assuming reads to follow a negative Binomial distribution with common dispersion parameter for a gene. This is implemented in grandR in the function *FitKineticsPulseR* using the code from Ref. [24] provided on github (https://github.com/dieterich-lab/ComparisonOfMetabolicLabeling).

### NTR method

For the Bayesian NTR method (method (iv) in the text), we note that under the assumption of steady state 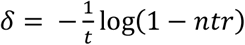. Thus, the posterior distribution of the NTR given data, *ntr*_*k*_ | *D*, can be transformed into a distribution on *δ*[15]. We assume *ntr*_*k*_ | ∼ *Beta* (*α*_*k*_, *β*_*k*_), and therefore the posterior density of the degradation rate is:

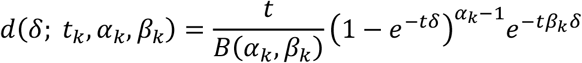

By logarithmizing and setting the derivative to 0, we see that the MAP estimator is

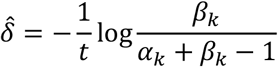

We can also transform the MAP of ntr_*k*_|*D*, yielding

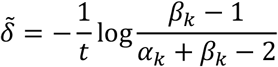

Thus, transforming from the *ntr* parameter to the degradation rate *δ* results in non-invariance of the MAP estimator. Both estimators are implemented in grandR, and we chose to use the transformed NTR MAP estimator 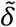 by default. With several samples, the degradation rate is estimated by numerically maximizing the log posterior

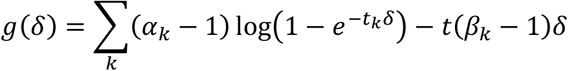

We use the *optimize* function built into R. For approximate *x*% credible intervals (CIs), we compute the critical drop in the log posterior distribution as 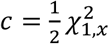 from a χ^2^ distribution with 1 degree of freedom similar to Ref. [24]. The rationale here is, that as we use a uniform prior, the posterior distribution is equal to the likelihood function. The CI is found by finding the values of *δ* left and right of the MAP estimate 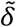, where 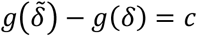. For exact CIs, we numerically integrate *g* using R’s *integrate* function and report the *x*% CI interval with the MAP as the central point. This is implemented in grandR’s function *FitKineticsGeneNtr*. It also provides an estimate of *σ* = *C*_*k*_ *δ*.

### Temporal recalibration

grandR implements two ways to recalibrate labeling times. The first can only be used with progressive labeling data and makes use of the fact that our kinetic model poses some constraints on how the temporal dynamics can behave. For that, we fit the NLLS model simultaneously for all genes, and consider the labeling time as additional variables that are jointly optimized. To make this procedure more efficient and less prone to noise, we first make a rough estimate of the half-lives using the uncalibrated labeling times and use the top 200 expressed genes from the following half-life classes: 0-2h,2-4h,6-8h, >8h. Stratifying by half-life classes is important as many of the most strongly expressed genes have very long RNA half-lives. Importantly, however, the labeling time parameters can only be estimated up to a constant factor which corresponds to the time unit of the model. We make this model identifiable by assuming that the effective labeling time is equal to the nominal labeling time for the last time point. This procedure is implemented in the function *CalibrateEffectiveLabelingTimeKineticFit* in grandR.

The second method for temporal recalibration requires reference half-lives. For each biological sample the observed data can be transformed into half-lives for any labeling time (see below, “Transforming snapshot data”). We choose the labeling time such that the median log fold change between the reference and transformed half-lives across all genes is 0 by using the *uniroot* function of R. This procedure is implemented in the function *CalibrateEffectiveLabelingTimeMatchHalflives* in grandR.

### Transforming snapshot data

By solving equations 5 and 6, we obtain

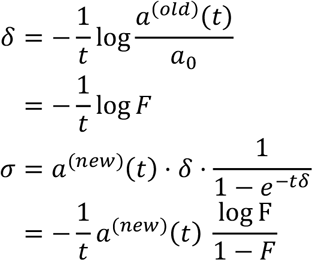

with 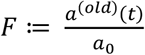. To compute *σ* and *δ* from this, the initial abundance *a*_0_ at time *t* = 0, i.e., at the start of labeling, must be known in addition to old and new RNA levels. This might not be the case, and only an abundance *a*′ at time *t*′ < 0 might be known, either by design of the experiments or because the effective labeling time is shorter than the nominal labeling time. In this case, the initial abundance can be computed as

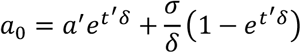

We use equation 6 to get rid of *σ*:

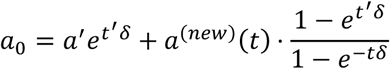

Substituting this into equation 5:

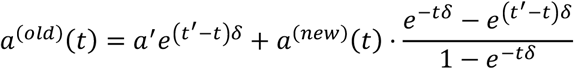

We solve this numerically for *δ* by using the R’s *uniroot* function. Of note, this assumes *σ* and *δ* to be constant throughout the time [*t* ′, *t*]. Transforming snapshot data is implemented in grandR’s *TransformSnapshot* function.

### Hierarchical Bayesian modeling of snapshot data

We define snapshot data for a single biological sample *k* and a single gene from a nucleotide conversion sequencing experiment to be a tuple 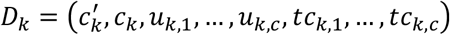. Here, 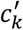 is the read count at the start of labeling at time *t* = 0, and *c*_*k*_ the read count at time *t*. For now, we ignore the need for normalization and assume that *c*_*k*_ and 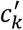 are directly comparable measures of gene expression, i.e. are already normalized. For ease of notation, we here assume a measurement *c*′ at *t* = 0, but we can adapt our model in principle also to situations, where the measurement is taken at any time *t*′ (see above, *Transforming snapshot data*). *u*_*k,r*_ and *tc*_*k,r*_ for *r*∈{1,…, *c*} represent the number of uridines and the number of T-to-C conversions, respectively, for a read *r*, i.e. the sufficient statistics for estimation of the *ntr* parameter. We will omit the index *k* if it is not necessary.

We assume that snapshot data *D* are generated by the following process:

1. Sample the unobserved parameters *a*_0_, *σ* and *δ* from unknown distributions representing the biological variability of the true initial abundance, the synthesis rate and degradation rate, respectively; this uniquely determines the full temporal kinetics of the true RNA abundance *a*(*t*) as well as *a*^(*new*)^(*t*) and *a*^(*old*)^(*t*) and 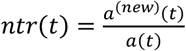
2. Sample *c* and *c* _0_ from unknown distributions with mean *a*(*t*) and *a*(0). These distributions represent the technical noise of the measurement.
3. Sample *u*_1_, …, *u*_*c*_ from the sequence of the gene. Which sequence is used depends on the protocol used for library preparation.
4. Sample *tc*_*r*_ for ∈ {1,…, *c*} from a Binomial mixture distribution

*BinomMix*(*u*_*r*_, *p*_*e*_, *p*_*c*_,*ntr* (*t*))

Here, we are mainly concerned with snapshot data 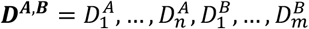 involving several biological replicates from two conditions *A* and *B*, and would like to infer the joint posterior distributions 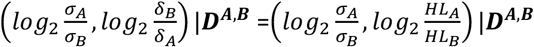. Note that for the synthesis rates we consider the log fold change *A* vs *B*, i.e. *B* is the *control* condition. We prefer to invert the log fold change of the degradation rates, which then corresponds to the more intuitive log fold change of the RNA half-lives *HL*_*A*_ vs *HL*_*B*_. Unfortunately, this is analytically intractable, and we found Markov chain Monte Carlo methods to be too inefficient considering the sheer size of *D*.

However, we show here that we can efficiently draw *N* samples (*σ*_1,_ *δ*_1_) … (*σ*_*N*_ *δ*_*N*_) from the joint posterior *σ, δ* | ***D*** for a single condition, with *n* replicate samples, i.e. ***D*** = *D*_1,_ … *D*_*n*_. Hence, 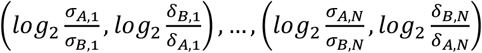 is a sample form the joint log fold change posterior distribution 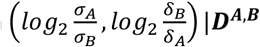.To draw a single sample (*σ*_*j*_, *δ*_*j*_) from the posterior *σ,δ*|***D***, we consider the following processes separately:

1. Draw a sample 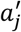 from the posterior distribution *a*_0_ | ***D*** = *a*(0) | ***D***.
2. Draw a sample *a*_*j*_ from the posterior distribution *a*(*t*) | ***D***.
3. Draw a sample *ntr*_*j*_ | ***D*** from the posterior distribution *ntr* (*t*) | ***D***.

We then transform these samples into *σ* and *δ* as described above under “Transforming snapshot data”. Note that the prior distribution for *σ,δ* as well as 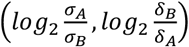 is thereby implicitly defined by the priors for *a*_0_, *a*(*t*), *ntr* (*t*).

### Sampling from *a*(.)| *D*

We assume that read counts *c*∼ *NegBinom* (*μ, d*) are distributed according to a negative Binomial distribution with mean *μ* and dispersion *d*. The dispersion parameter is defined above such that the variance is *μ* + *dμ* ^2^. To enable efficient sampling, we assume *d* to be fixed (for a single gene) and use *estimateDispersions* from the DESeq2 package for estimation. There is no obvious conjugate prior for *μ*, however, we can reparametrize the negative Binomial *NegBinom*′ (*s, p*) by 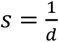 and 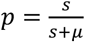.Then, 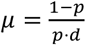

It is easy to see that the Beta distribution is a conjugate prior for *p*: Given *n* samples ***c*** = *c*_1_, …, *c*_*n*_, the density of the posterior for *p* for a *NegBinom*′ (*s, p*)likelihood and *Beta*(*α, β*) prior is

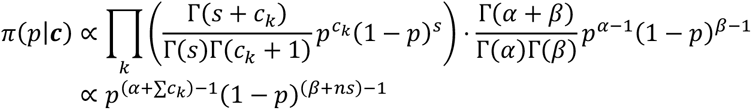

Thus, for the prior *p*∼ *Beta*(*α β*), we have the posterior *p* | *c*_1,_ …, *c*_*n*_ ∼ *Beta*(*α*+ ∑*c*_*k*_, *β* + *ns*).

We use the full distribution of all genes to inform the prior distribution as follows. We first transform the expression value *c*_*i*_ for each gene *i* to 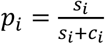 with 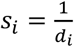 and use the method of moments to fit the hyperparameters *α* and *β*, which we then use for the whole data set of all genes.

So far, we have ignored normalization. For practical applications, this must be taken into account. We do this by the same approach as DESeq2, i.e. by rescaling read counts using a size factor to obtain normalized read counts [20]. This can be achieved in grandR by first calling the *Normalize* function, which places the normalized read counts into the default *data slot* of the grandR object.

Thus, to sample from *a*(*t*) | *D*_1_,…, *D*_*n*_, we draw random numbers from a *Beta* 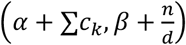 to sample from *a*_0_| *D*_1_…, *D*_*n*_, we draw random numbers from a *Beta* 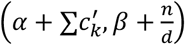 distribution. Here, *c*_*k*_ and 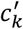 are the normalized read count from time *t* and 0, respectively, of data set *D*_*k*_, *d* is the dispersion parameter estimated by DESeq2, and *α* and *β* are the prior hyperparameters. Each of these Beta distributed values *p* is then transformed via 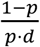 to obtain a sample from *a*(*t*) | *D*_1_…, *D*_*n*_ or *a*_0_| *D*_1_…, *D*_*n*_.

### Sampling from *ntr* (*t*) | ***D***

The number of conversions on a read *tc*_*r*_∼ *BinomMix*(*u*_*r*_, *p*_*e*_, *p*_*c*_, *ntr*) are distributed according to a Binomial mixture distribution as defined above. The number of uridines *u*_*r*_ is fixed, and to enable efficient sampling, we also assume the parameters *p*_*e*_ and *p*_*c*_ to be fixed. The posterior distribution *ntr*| *tc*_1_,…, *tc*_*r*_ for a single biological sample, which is computed numerically by GRAND-SLAM, can be approximated by a Beta distribution, and we assume this Beta to be conjugate with the Beta prior used by GRAND-SLAM to compute the posterior distribution [15]. This posterior only quantifies technical variance of measuring the true *ntr* for a single biological sample. To handle biological variability in addition, we introduce an additional hierarchical layer in our Bayesian model:

For each biological sample *k* ∈{1,…, *n*}, we have *ntr*_*k*_ | ***D***_*k*_∼ *Beta*(*α* + *α*_*k*_, *β* + *β*_*k*_). Here, *α* and *β* are the parameters of the prior Beta distribution reflecting biological variability of *ntr* across biological replicate samples and *α*_*k*_ and *β*_*k*_ are the parameters estimated by GRAND-SLAM from the given *tc*_*k*,1_, …, *tc*_*k,r*_, which reflect technical noise. The joint density of all ***ntr*** = (*ntr*_1_,…, *ntr*_*n*_) is

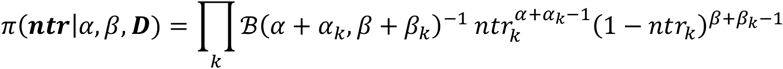

Here, ℬ is the beta function. When imposing a prior on (*α, β*), the joint posterior of all parameters is

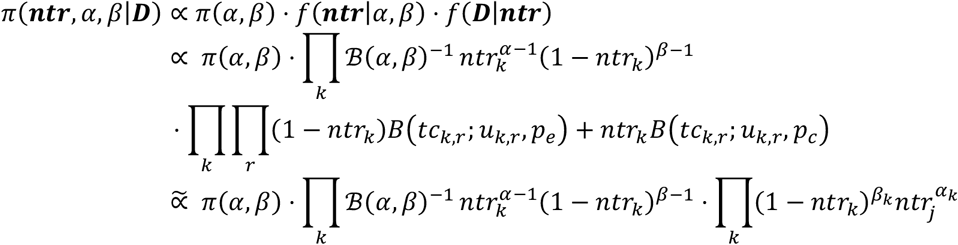

Here, 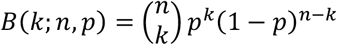, and the last line follows from our Beta approximation of the mixture model. Thus, the marginal posterior distribution of (*α, β*) is

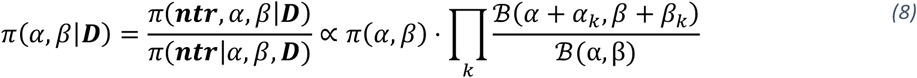

If the marginal posteriors *ntr*_*k*_|***D***_***k***_ overlap significantly, a point *ntr* and, therefore, a *Beta* (*α, β*) prior with infinitesimally small variance or, equivalently, infinite *α* + *β* becomes probable. An appropriate constraint can be imposed using the prior distribution *π* (*α, β*). We decided to use the following sigmoid function

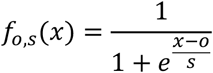

This can be integrated:

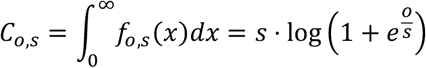

and thus,

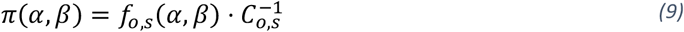

is a proper prior. *f*_*o,s*_ is almost constant before the offset *o* and quickly (depending on *s*) goes to zero after *o*, i.e. *o* represents a maximal *α* + *β*, or, equivalently, minimal variance, that has substantial prior probability. We set *o* such that the variance of the prior *π* (*α, β*) is equal to the sample variance 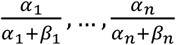. Importantly, as long as (i) the mean 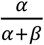 is unconstrained and (ii) the minimal variance is constrained, the exact choice of the prior *π* (*α, β*) only has minor effect on sampling of *ntr*| ***D***.

To sample *ntr*|***D***, i.e. the mean 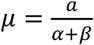 from the distribution *π*(*α, β*| ***D***;), we compute the marginal posterior on a grid of values [30]. Since we want to sample *μ*, it makes sense not to build an (*α, β*) - grid, but to reparametrize and build the grid with coordinates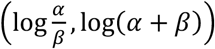 [30]. Note that 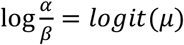. For each grid point (*x, y*), we transform 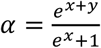 and 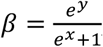, for which we compute the unnormalized posterior density defined in equation 8 with prior from equation 9, and, due to our reparametrization, multiply this by the Jacobian determinant

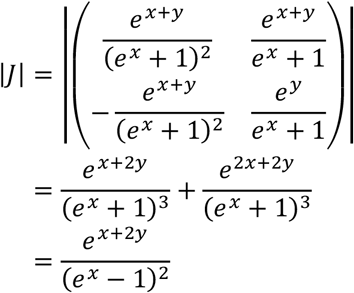

For numerical stability, we compute everything in log space, then subtract the maximal grid value and exponentiate [30]. To determine the grid bounds, we first find the maximum using R’s *optim* function, and then go into positive and negative *x* and *y* directions to see where the grid would drop below 1000-fold of the maximal value using R’ *uniroot* function. To sample *μ*, we first sum over the columns of the grid and normalize to obtain a discrete probability distribution *l*_1_, … *l*_*m*_. Note that each *l*_*j*_ corresponds to a particular value of 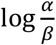. One of these values is sampled from the distribution *l*_1_, … *l*_*m*_ and random uniform jitter is added to fill the spacing of the grid [30]. This value *x* is then transformed to *μ* = *logit*^−1^(*x*).

## Availability of Data and Materials

Project name: grandR

Package version: 0.2.0

Project home page: https://github.com/erhard-lab/grandR

Archived version: https://CRAN.R-project.org/package=grandR

Operating system(s): Platform independent

Programming language: R

License: Apache License 2.0

Raw data sets used here are available at GEO under accession numbers GSE99970 (24h 4sU labeling data), GSE139151 (NXF1 knockdown data), GSE162323 (SARS-CoV-2 data), and GSE155604 (BANP depletion data). All processed data (GRAND-SLAM outputs) are available on zenodo under https://doi.org/10.5281/zenodo.6513333 (24h 4sU labeling data), https://doi.org/10.5281/zenodo.5907183 (NXF1 knockdown data), https://doi.org/10.5281/zenodo.5834034 (SARS-CoV-2 data), and https://doi.org/10.5281/zenodo.6976391 (BANP depletion data).

R notebooks and data files for generating all figures are available on https://github.com/erhard-lab/grandR-manuscript/releases/tag/init.submission.

## Author information

### Contributions

All authors performed the computational analyses and produced the figures for the manuscript, FE wrote the source codes for the grandR package, and FE wrote the manuscript with contributions of all authors. All authors read and approved the final manuscript.

### Authors’ information

Twitter handle: @erhard_lab

### Corresponding authors

Correspondence to Florian Erhard

### Ethics declarations

Not applicable.

### Ethics approval and consent to participate

Not applicable.

## Figure legends

**Figure S1:**
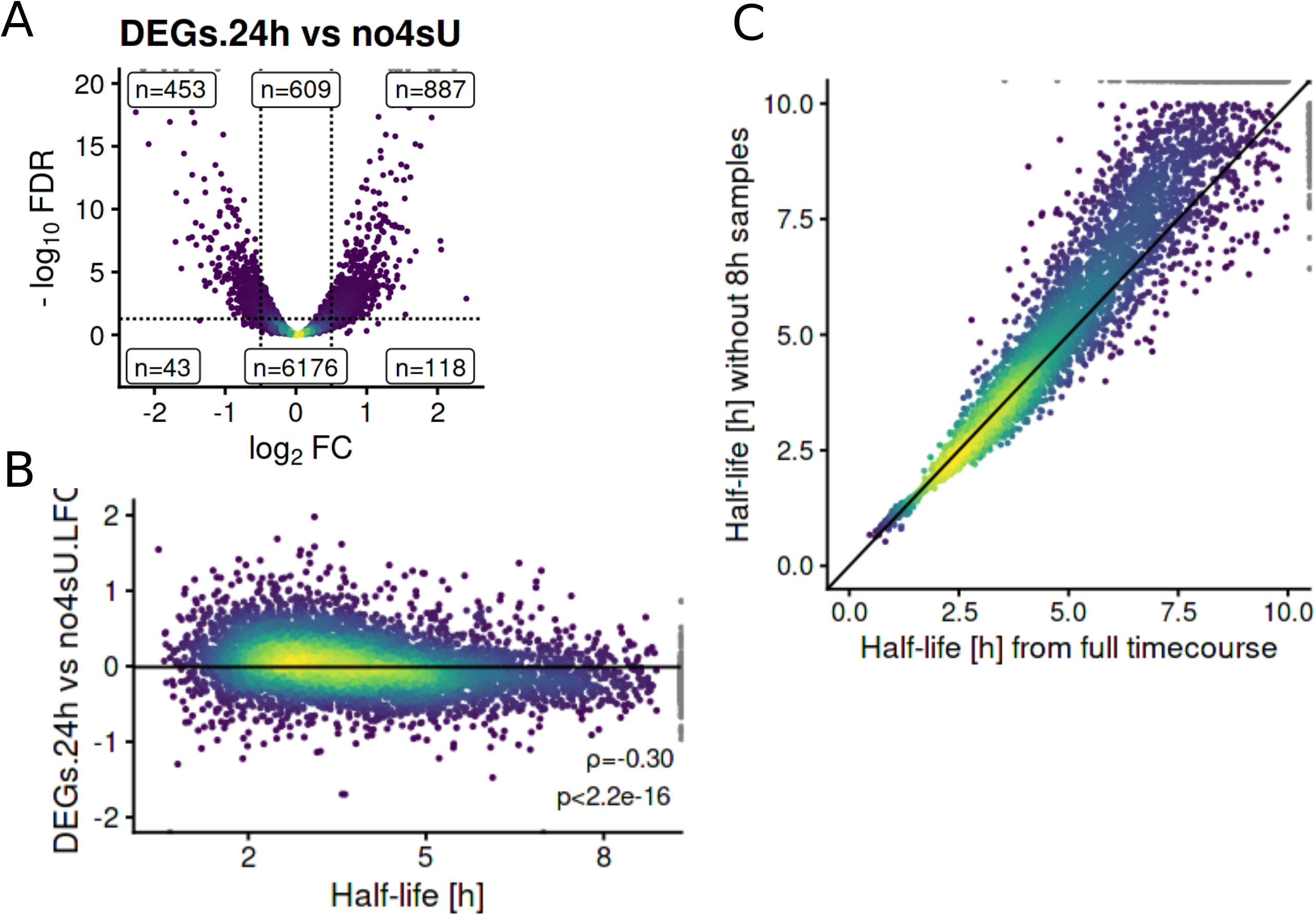
Quality control using grandR. **A** Vulcano plot of differentially expressed genes on total RNA level. The y axis shows the DESeq2 P value (Wald test) adjusted for multiple testing (Benjamini-Hochberg; FDR, false discovery rate). The numbers of genes above and below 5% FDR and for a threshold of 1.4-fold up- or downregulation are indicated. **B** RNA half-lives (taken from Ref. [6]) are scattered against the log2 fold change of the 24h samples vs. control samples. Spearman correlation coefficient with associated P value (approximate t test) is indicated. **C** Scatterplot showing half-lives computed using the non-linear least squares method for the untreated mock samples from Ref. [18] for each gene. The x axis shows the half-life values considering the full time course (0h,2h,4h,8h), whereas the y axis shows the half-life values after excluding the 8h time point.

**Figure S2:**
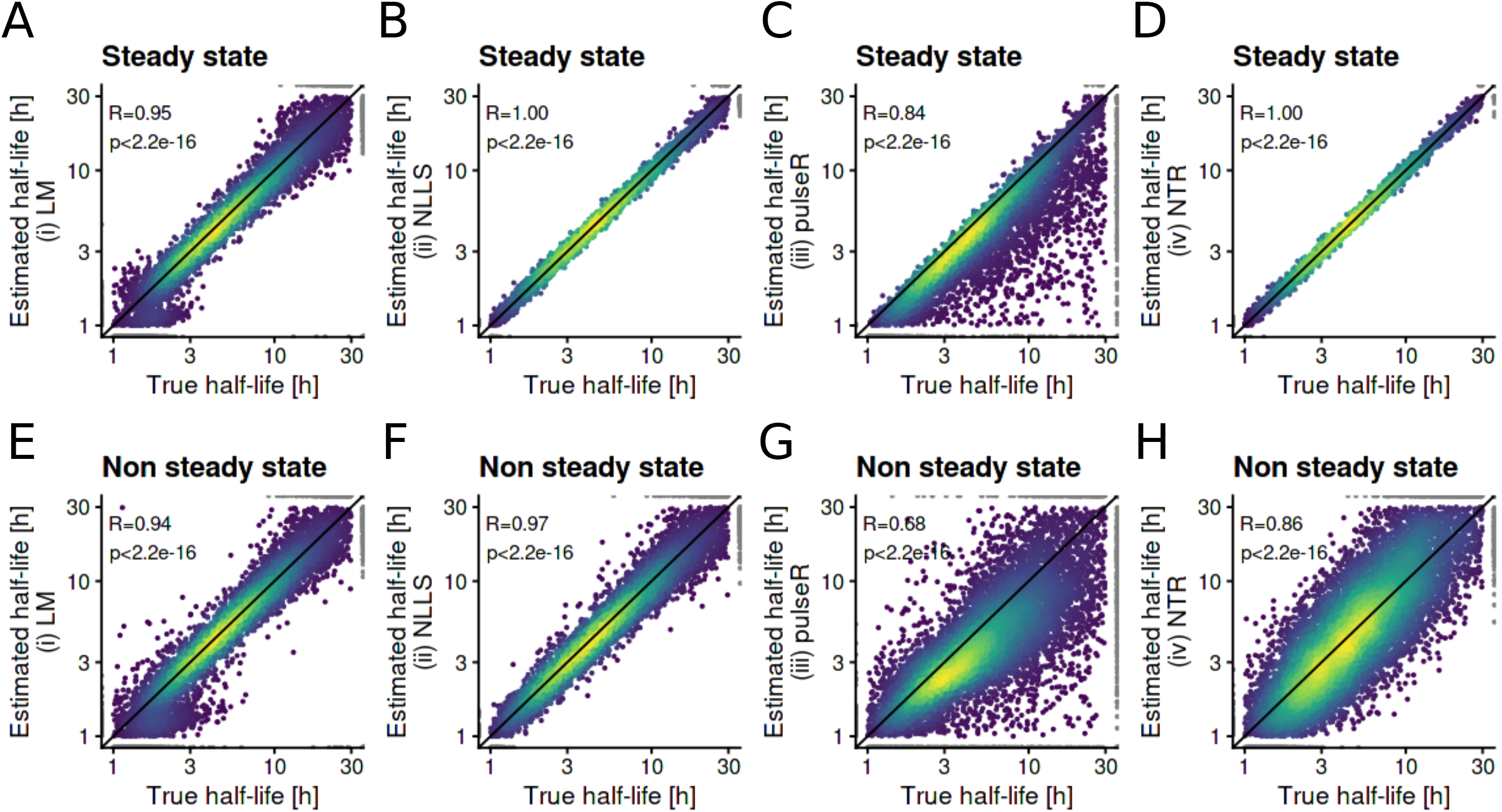
Evaluation of half-life estimates from simulated data (0h,1h,2h,4h,8h, 3 replicates each, 20 million reads per sample; half-lives and expression values were taken from the Mock samples of Ref. [17]). Scatterplots compare the true, simulated half-life value for each gene against the half-life value estimated by the linear model (**A** and **E**), the non-linear least squares approach (**B** and **F**), the pulseR method (**C** and **G**) and the Bayesian approach (**D** and **H**). Results for simulated steady state gene expression (**A-D**) and when gene expression at 0h was perturbed (**E-H**) are shown. The Pearson correlation coefficient and associated P values (approximate t test) are indicated.

**Figure S3:**
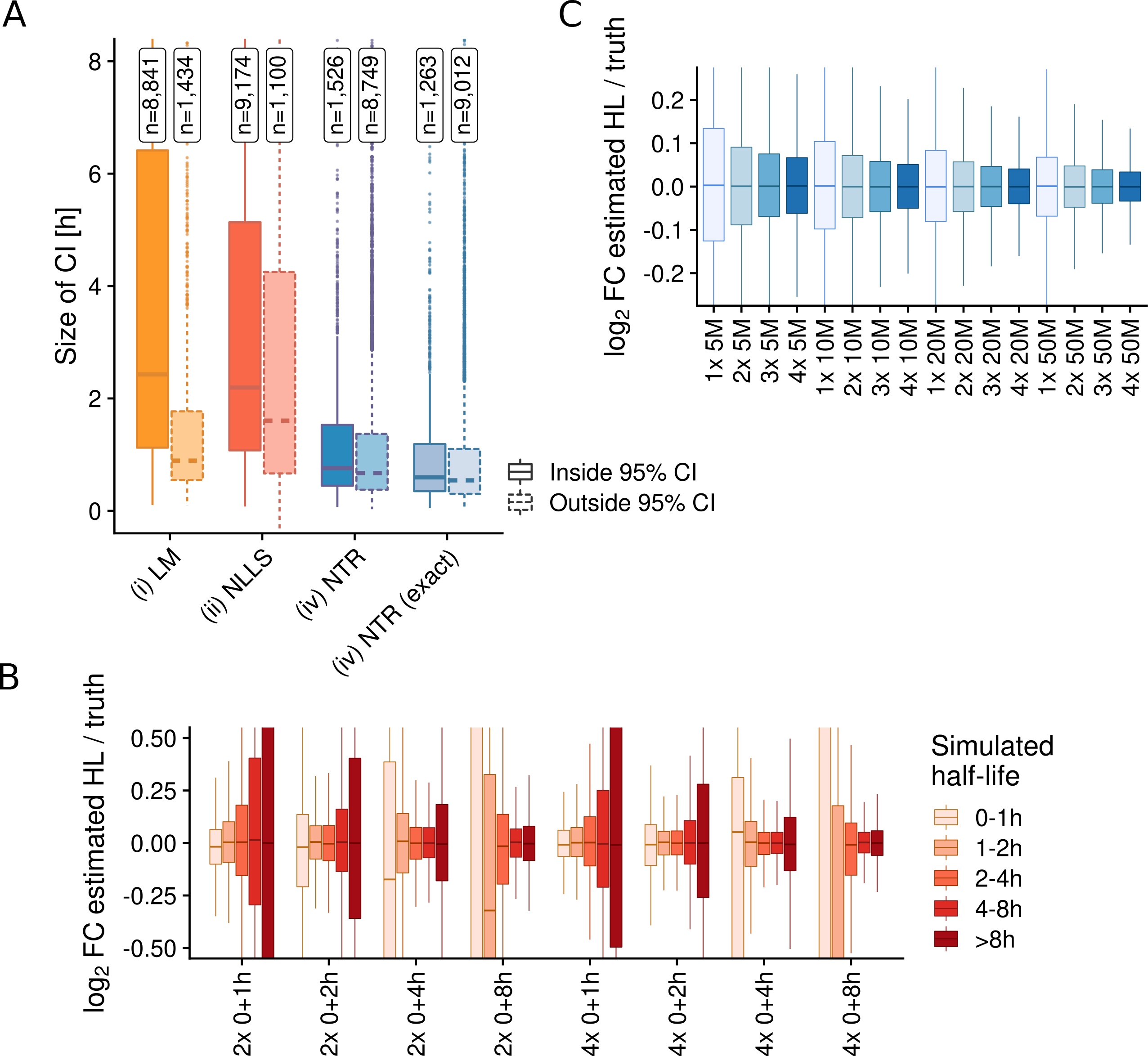
Estimating half-lives using progressive labeling experiments. **A** Boxplots showing the sizes of 95% half-life confidence intervals (CI; for LM and NLLS) or 95% half-life credible intervals (CI; for NTR). Simulations were performed under steady state conditions where the initial value was *a*_0_ ≠ *σ*/*δ* for each gene. Distributions for genes having the ground-truth inside or outside of the estimated CI are shown separately and the numbers of these genes are indicated. NTR represents the χ^2^ approximation of CIs, NTR (exact) represents exact CIs computed numerically. **B** Boxplots showing log2 fold changes of half-lives estimated by the NLLS method vs the ground truth of simulated data under steady state conditions. The distributions for different half-life classes are shown for several experimental settings involving the indicated number of replicates and time points. **C** Boxplots showing log2 fold changes of half-lives estimated by the NLLS method vs the ground truth of simulated data under steady state conditions for a full progressive labeling time course (1h,2h,4h and 8h). The distributions involving the indicated number of replicates and sequencing depth in million (M) reads are shown.

**Figure S4:**
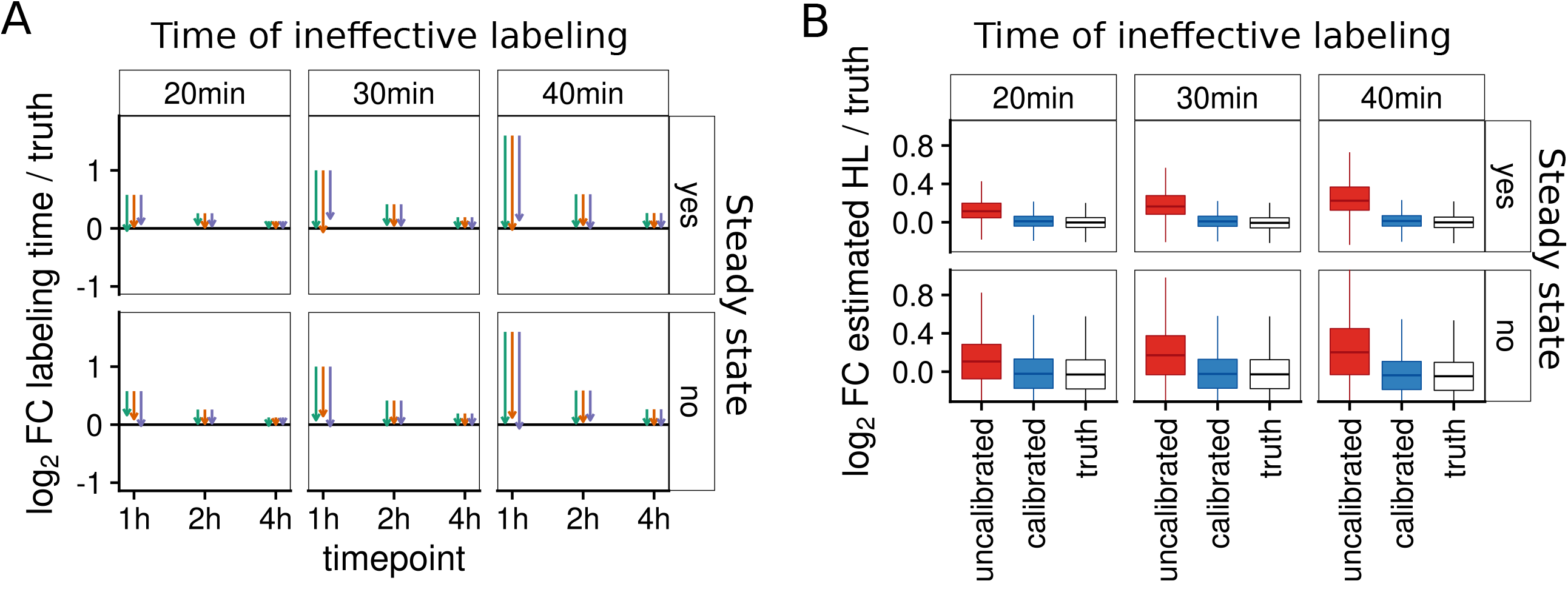
Temporal recalibration of simulated data. **A** Plot comparing labeling times before and after recalibration for simulated data. Data was simulated based on SARS-CoV2-data (nominal labeling times 0h,1h,2h,4h,8h) either under steady state conditions or non-steady state conditions as indicated. For the 1-4h time points, an ineffective time of labeling as indicated was subtracted from the nominal times before simulation. Arrows show the log2 fold change of uncalibrated, nominal labeling times vs the true effective labeling time (start of the arrow) and of the recalibrated labeling time vs the true effective labeling time (tip of the arrow). Three replicates are shown by colors. **B** Boxplots showing log2 fold changes of estimated (NLLS) half-lives vs. ground truth for n=9,162 genes before recalibration (uncalibrated), after recalibration (calibrated) and when the true effective labeling times were used (truth).

**Figure S5:**
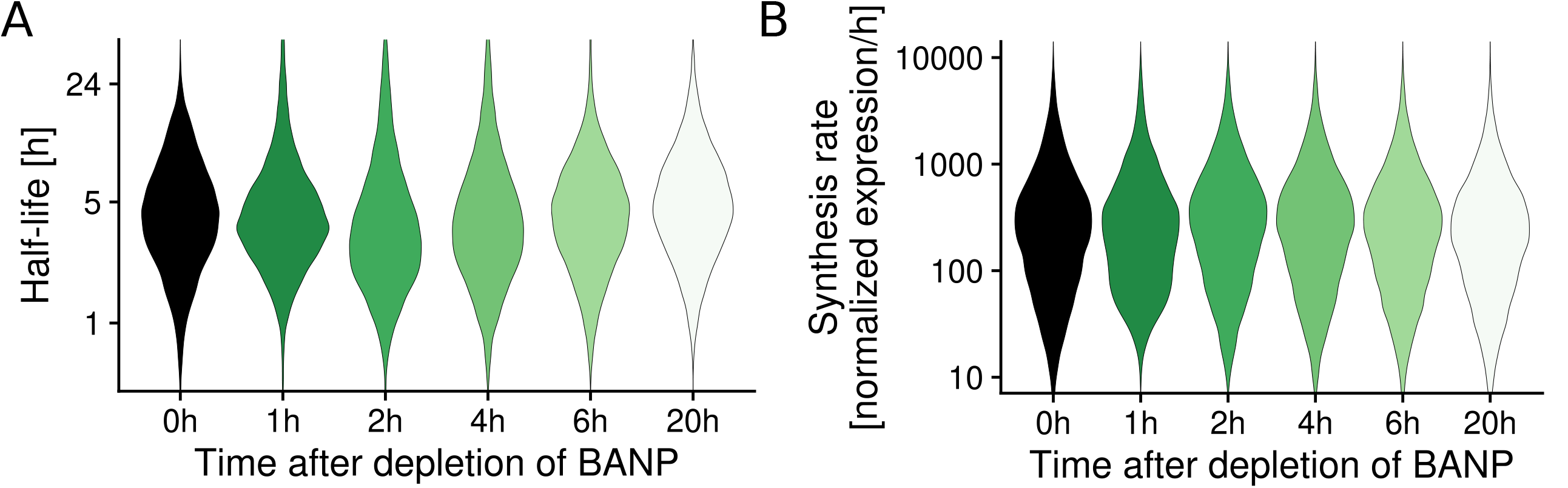
Temporal recalibration of BANP depletion experiments. Violin plots showing the distribution of estimated RNA half-lives (**A**) and synthesis rates (**B**) for n=11,096 genes estimated by our Bayesian hierarchical model for each experimental time point after temporal recalibration.

