## Supplementary data file 1 for "grandR: a comprehensive package for nucleotide conversion sequencing data analysis": Fig1.Rmd

````
---
title: "Figure 1"
output:
  html_document:
  dev: "svg"
---


We load the GRAND-SLAM output and preprocess as usual. Then we fit the kinetic model and present data in a web-interface, using the PlotGeneKinetics function for visualizing individual genes.
```{r eval=1:8}
suppressPackageStartupMessages({
    library(grandR)
    library(ggplot2)
    library(patchwork)
})
file <- "https://zenodo.org/record/5834034/files/sars.tsv.gz"
desc <- c(Design$Condition,Design$dur.4sU,Design$Replicate)
sars <- ReadGRAND(file,design=desc)
sars <- FilterGenes(sars)
sars <- Normalize(sars)
stst <- c(Mock=TRUE,SARS=FALSE)
sars <- FitKinetics(sars,steady.state=stst)
ServeGrandR(sars,plot.gene = Defer(PlotGeneProgressiveTimecourse,steady.state=stst))
```
````
