## Supplementary data file 1 for "grandR: a comprehensive package for nucleotide conversion sequencing data analysis": Fig2A-B_S1.Rmd

````
---
title: "Figure 2A-B, S1"
output: 
  html_document:
    dev: "svg"
    keep_md: yes
---

We first load the GRAND-SLAM output and preprocess as usual.
```{r}
suppressPackageStartupMessages({
    library(grandR)
    library(ggplot2)
    library(patchwork)
})

d <- ReadGRAND("https://zenodo.org/record/6513333/files/ameres_24h.tsv.gz",design=c("Cell",'4sU',Design$Replicate))
d <- FilterGenes(d)
```

Now, we compute a principal component analysis (PCA). The PCA is by default computed from the total count matrix after 
variance stabilizing transformation using the vst function of the DESeq2 package, and using the top 500 variable genes.

```{r Fig2A, fig.height=2.5, fig.width=4}
PlotPCA(d,aest = aes(color=`4sU`,shape=NULL))+
  scale_color_manual(values=c(no4sU='gray50',`24h`='#f03b20'))
```

Compute differential gene expression of total RNA using the Wald test of DESeq2 (Benjamini-Hochberg multiple testing adjusted), 
and log2 fold changes using the lfc package. Show the vulcano plot:

```{r FigS1A, fig.height=3, fig.width=3, dev='CairoPNG'}
contr <- GetContrasts(d,contrast = c("4sU","no4sU"),no4sU = TRUE)
d <- PairwiseDESeq2(d,name="DEGs",contrasts=contr,mode="total")
d <- LFC(d,contrast=contr,name="DEGs",mode="total")
VulcanoPlot(d,lfc.cutoff = 0.5,ylim=c(-3,20))
```


Perform GSEA on MsigDB Hallmark gene sets. AnalyzeGeneSets returns all pathways with Benjamini-Hochberg multiple testing adjusted P values < 5%.
```{r Fig2B, fig.height=3.5, fig.width=6}
gsea.res <- AnalyzeGeneSets(d,criteria = LFC, category="H",process.genesets = function(n) gsub("HALLMARK_","",n),verbose=FALSE)
enrichplot::ridgeplot(gsea.res)
```

Download the suppementary Excel table containing RNA half-lives from Herzog et al., and plot these against the log2FC between 24h 4sU treatment and no4sU control (and compute Spearman correlation):
```{r FigS1B, fig.height=2.7, fig.width=4, dev='CairoPNG'}
library(readxl)
tmp <- tempfile(fileext = "xls")
download.file(url = "https://static-content.springer.com/esm/art%3A10.1038%2Fnmeth.4435/MediaObjects/41592_2017_BFnmeth4435_MOESM4_ESM.xls", destfile = tmp)
tab <- read_excel(tmp)
file.remove(tmp)

df <- GetAnalysisTable(d,columns="LFC",genes = intersect(Genes(d),tab$Name))
df$`Half-life [h]`=setNames(tab$`Half-life (h)`,tab$Name)[rownames(df)]
PlotScatter(df,x=`Half-life [h]`,y=`DEGs.24h vs no4sU.LFC`,ylim=c(-2,2),correlation = FormatCorrelation(method="spearman"),correlation.x = Inf,correlation.y = -Inf)+geom_hline(yintercept = 0)+scale_x_continuous(breaks=c(2,5,8))
```
````
