## Supplementary data file 1 for "grandR: a comprehensive package for nucleotide conversion sequencing data analysis": Fig2C_S1C.Rmd

````
---
title: "Figure 2C, S1C"
output: 
  html_document:
    dev: "svg"
    keep_md: yes
---

We first load the GRAND-SLAM output and preprocess as usual.
```{r}
suppressPackageStartupMessages({
    library(grandR)
    library(ggplot2)
    library(patchwork)
})

d <- ReadGRAND("https://zenodo.org/record/5907183/files/nxf1.tsv.gz",design=c("Cell",'Cyto',Design$dur.4sU,Design$has.4sU, Design$Replicate))
d <- FilterGenes(d)
d <- Normalize(d)
```

Now, generate the toxicity control plots for replicates A of 2h, 4h, and 8h:
```{r Fig2C, dev='CairoPNG', fig.height=3, fig.width=5}
PlotToxicityTestRank(d,w4sU = "mock.cyto.2h.4sU.A",ylim = c(-2,2))
PlotToxicityTestRank(d,w4sU = "mock.cyto.4h.4sU.A",ylim = c(-2,2))
PlotToxicityTestRank(d,w4sU = "mock.cyto.8h.4sU.A",ylim = c(-2,2))
```


To check, we fit the kinetic model with and without the 8h timepoint, and compare the fitted Half-lives.

```{r FigS1C, dev='CairoPNG', fig.width=4, fig.height=4}
d <- subset(d,columns = Cell=="mock")
SetParallel()
d <- FitKinetics(d,name="0-8")
d <- subset(d,duration.4sU<8)
d <- FitKinetics(d,name="0-4")
df <- GetAnalysisTable(d,columns="Half-life")
PlotScatter(df,x=`0-8.Half-life`,y=`0-4.Half-life`,xlim=c(0,10),ylim=c(0,10))+
  geom_abline()+
  xlab("Half-life [h] from full timecourse")+
  ylab("Half-life [h] without 8h samples")
```
````
