## Supplementary data file 1 for "grandR: a comprehensive package for nucleotide conversion sequencing data analysis": Fig3A-C_S2S3A.Rmd

````
---
title: "Figure 3A-C and S2, S3A"
output: 
  html_document:
    dev: "svg"
    keep_md: yes
---


Load default setting data (3 replicates, 1,2,4,8 h, 20M reads) 
To see how this data was generated have a look into the loadGenerate R script.

```{r}
suppressPackageStartupMessages({
    library(grandR)
    library(ggplot2)
    library(patchwork)
})

source("loadGenerate.R")
data_default=loadGenerate(ProgressiveLabelingTimecourseAllModelsFitted)

# define labels and colors
methods=c(`(i) LM`="lm.ss.Half-life",`(ii) NLLS`="nlls.ss.Half-life",`(iii) pulseR`="pulseR.ss.Half-life",`(iv) NTR`="ntr.ss.Half-life")
methods.ci=c(`(i) LM`="lm.ss.Half-life",`(ii) NLLS`="nlls.ss.Half-life",`(iv) NTR`="ntr.ss.Half-life",`(iv) NTR (exact)`="entr.ss.Half-life")
method.colors=c(`(i) LM`="#fe9929",`(ii) NLLS`="#fb6a4a",`(iii) pulseR`="#756bb1",`(iv) NTR`="#2b8cbe",`(iv) NTR (exact)`="#a6bddb")
```

Scatter the true (simulated) RNA half-lives vs. the half-lives inferred for each method. Do this for both steady state (ss) and non steady state(noss) conditions.

```{r FigS2A-H, fig.height=3, fig.width=3, warning=FALSE, dev='CairoPNG'}
for (ss in c("ss","noss")) for (y in names(methods)) {
  print(
    PlotScatter(data_default,x=log(2)/true_d,ycol=gsub("ss",ss,methods[y]),xlim=c(1,30),ylim=c(1,30),log=TRUE,correlation = FormatCorrelation())+
      geom_abline()+
      xlab("True half-life [h]")+
      ylab(paste0("Estimated half-life [h]\n",y))+
      ggtitle(if(ss=='ss') "Steady state" else "Non steady state")
  )
}
```
Compute the log2 fold change of half-lives (true vs estimated) for all methods and show them in an empirical cumulative distribution plot. Do this for both steady state (ss) and non steady state(noss) conditions.

```{r Fig3A-B, fig.height=2.5, fig.width=4.5, warning=FALSE}
pp=function(pattern) {
  tab = log2(as.matrix(GetAnalysisTable(data_default,analyses = pattern,columns='Half-life$',gene.info = FALSE)*GeneInfo(data_default)$true_d/log(2)))
  colnames(tab)=gsub("noss","ss",colnames(tab))
  tab = tab[,methods]
  colnames(tab)=names(methods)
  tab = reshape2::melt(tab, id.vars =c(),varnames = c("Gene","Method"),value.name = "logFC")
  ggplot(tab, aes(x=logFC, color=Method)) + 
    cowplot::theme_cowplot()+
    geom_vline(xintercept = 0, linetype=2,color='gray')+
    geom_hline(yintercept = 0.5, linetype=2,color='gray')+
    stat_ecdf(geom = "line",size=0.8)+ 
    coord_cartesian(xlim=c(-1.5,1.5)) + 
    xlab(bquote(log[2]~FC)) + 
    scale_color_manual(NULL,values=method.colors)+
    scale_y_continuous('Cumulative frequency',breaks=c(0,0.5,1),labels = c("0","0.5","1"))
}

pp("\\.ss")+ggtitle("Steady state")
pp("\\.noss")+ggtitle("Non steady state")
  
```

Analyze the size of the confidence or credible intervals

```{r Fig3C-FigS3A, fig.height=5, fig.width=5, warning=FALSE}
pp=function(pattern) {
  tab = GetAnalysisTable(data_default,analyses = pattern, columns = "Half-life", gene.info = FALSE)
  colnames(tab)=gsub("noss","ss",colnames(tab))
  tab$true_HL=log(2)/GeneInfo(data_default)$true_d
  tab=tab[tab$true_HL<24,]
  make.df=function(method) {
    data.frame(
      HL=cut(tab$true_HL,c(0,2,4,8,Inf)),
      CI=tab[[paste0(methods.ci[method],".upper")]]-tab[[paste0(methods.ci[method],".lower")]],
      inside=ifelse(tab$true_HL>tab[[paste0(methods.ci[method],".lower")]] & tab$true_HL<tab[[paste0(methods.ci[method],".upper")]],'Inside 95% CI',"Outside 95% CI"),
      method=method
      )
  }
  df=do.call("rbind",lapply(names(methods.ci),make.df))
  df2=as.data.frame(table(df[,c("method","inside")]))
  ggplot(df,aes(method,CI,fill=method,color=method,linetype=inside,alpha=inside))+
    cowplot::theme_cowplot()+
    geom_boxplot(outlier.size = 0.1, outlier.alpha = 0.5)+
    scale_fill_manual(NULL,values=method.colors,guide="none")+
    scale_color_manual(NULL,values=colorspace::darken(method.colors,0.1),guide="none")+
    scale_alpha_manual(values=c(1,0.5),guide="none")+
    coord_cartesian(ylim=c(0,8))+
    ylab("Size of CI [h]")+xlab(NULL)+
    scale_linetype(NULL)+
    ggtext::geom_richtext(data = df2,aes(method,Inf,label=sprintf("n=%s",scales::label_comma()(Freq)),group=inside),alpha=1,color='black',fill='#FFFFFFB3',hjust=1.1,angle=90,position = position_dodge(width = .75),label.padding = unit(c(0.1, 0.25, 0.1, 0.25), "lines"))+
    theme(axis.text.x = element_text(angle = 45, vjust = 1, hjust=1))
}

pp("\\.ss")+ggtitle("Steady state")
pp("\\.noss")+ggtitle("Non steady state")
ggplot_build(pp("\\.ss"))$data[[1]][,1:9]
```
````
