## Supplementary data file 1 for "grandR: a comprehensive package for nucleotide conversion sequencing data analysis": Fig3D_S3B-C.Rmd

````
---
title: "Figure 3D, S3B-C"
output: 
  html_document:
    dev: "svg"
    keep_md: yes
---


```{r}
suppressPackageStartupMessages({
    library(grandR)
    library(ggplot2)
    library(patchwork)
})

source("loadGenerate.R")
data=loadGenerate(ProgressiveLabelingSettings)
```

```{r Fig3D,fig.width=8,fig.height=3}
pp=function(cols) {
  tab = GetAnalysisTable(data,columns=c('Half-life$'),gene.info = FALSE)[,cols]
  names(tab)=gsub(".Half-life","",names(tab))
  tab=as.data.frame(lapply(tab,function(hl) log2(hl*GeneInfo(data)$true_d/log(2))),check.names=FALSE)
  tab$HL=log(2)/GeneInfo(data)$true_d
  
  tab=reshape2::melt(tab,id.vars = "HL",variable.name = "Setting",value.name = "logFC")
  tab$Setting=factor(tab$Setting,levels=Analyses(data))
  tab$HL.cat=cut(tab$HL,breaks=c(0,1,2,4,8,Inf),labels = c("0-1h","1-2h","2-4h","4-8h",">8h"))
  
  ggplot(tab, aes(Setting,y=logFC,fill=HL.cat,color=HL.cat)) + 
    cowplot::theme_cowplot()+
    geom_boxplot(outlier.shape = NA,size=0.2)+
    coord_cartesian(ylim = c(-0.5,0.5))+
    theme(axis.text.x = element_text(angle=90, hjust = 1, vjust = 0.5)) + 
    xlab(NULL) + 
    ylab(bquote(log[2]~FC~"estimated HL / truth")) +
    scale_fill_manual("Simulated\nhalf-life",values=RColorBrewer::brewer.pal(5,'Reds'))+
    scale_color_manual("Simulated\nhalf-life",values=colorspace::darken(RColorBrewer::brewer.pal(5,'Reds'),0.4))
}
pp(9:25)

```

```{r FigS3B,fig.width=7,fig.height=2.5}
pp(1:8)
```

```{r FigS3C,fig.width=4,fig.height=2.5}
data=loadGenerate(ProgressiveLabelingDepth)


tab = GetAnalysisTable(data,columns=c('Half-life$'),gene.info = FALSE)
names(tab)=gsub("(\\d+)M.(\\d+)R.Half-life","\\2x \\1M",names(tab))
lvl=names(tab)

tab=as.data.frame(lapply(tab,function(hl) log2(hl*GeneInfo(data)$true_d/log(2))),check.names=FALSE)
tab=reshape2::melt(tab,id.vars = c(),variable.name = "Setting",value.name = "logFC")
tab$Setting=factor(tab$Setting,levels=lvl)

ggplot(tab, aes(Setting,y=logFC,color=Setting,fill=Setting)) + 
    cowplot::theme_cowplot()+
  geom_boxplot(outlier.shape = NA,size=0.2)+
  coord_cartesian(ylim = c(-0.25,0.25))+
  theme(axis.text.x = element_text(angle=90, hjust = 1, vjust = 0.5)) + 
  xlab(NULL) + 
  ylab(bquote(log[2]~FC~"estimated HL / truth")) +
  scale_fill_manual(NULL,values=rep(RColorBrewer::brewer.pal(4,'Blues'),4),guide="none")+
  scale_color_manual(NULL,values=rep(colorspace::darken(RColorBrewer::brewer.pal(4,'Blues'),0.4),4),guide="none")

```
````
