## Supplementary data file 1 for "grandR: a comprehensive package for nucleotide conversion sequencing data analysis": Fig4A-D.Rmd

````
---
title: "Figure 4A,B,E,F"
output: 
  html_document:
    dev: "svg"
    keep_md: yes
---

Load recalibration data (SARS data, contains already the recalibrated times in Coldata in addition to the nominal times) 
To see how this data was generated have a look into the loadGenerate R script.


```{r}
suppressPackageStartupMessages({
    library(grandR)
    library(ggplot2)
    library(patchwork)
})
source("loadGenerate.R")

d=loadGenerate(CalibrateSARS)
```

Plot SMAD3 as an example, before and after recalibration

```{r Fig4A, fig.width=4,fig.height=2.3}
m=subset(d,Condition=='Mock')
Condition(m)=NULL
PlotGeneProgressiveTimecourse(m,"SMAD3",steady.state=list(Mock=TRUE,SARS=FALSE))
PlotGeneProgressiveTimecourse(m,"SMAD3",steady.state=list(Mock=TRUE,SARS=FALSE),time = "calibrated_time",exact.tics = FALSE)
```

Plot the distributions of the relative residuals (i.e. the residual divided by the model fit value) for each time point.

```{r Fig4B_left, fig.width=3,fig.height=2.4}
SetParallel()
m <- subset(d,columns=Condition=="Mock")
m <- FitKinetics(m,type="nlls",compute.residuals=TRUE,return.extra = function(s) setNames(s$residuals$Relative,paste0("Residuals.",s$residuals$Name))[s$residuals$Type=="new"])

df <- reshape2::melt(GetAnalysisTable(m,columns = "Residuals", gene.info = FALSE)[,-1], id.vars=c(),variable.name = "Sample", value.name = "Residual")
df$Sample=plyr::revalue(df$Sample,setNames(sub("kinetics.Mock.Residuals.","",levels(df$Sample)),levels(df$Sample)))
ggplot(df,aes(Sample,Residual,fill=Sample))+
    cowplot::theme_cowplot()+
  geom_hline(yintercept = 0, linetype = 2)+
  geom_boxplot(outlier.size = 0.1)+
  scale_fill_brewer(palette="Reds",guide="none")+
  coord_cartesian(ylim=c(-1,1))+
  scale_y_continuous("Relative residual",labels = scales::percent_format(1))+
  theme(axis.text.x = element_text(angle = 45, vjust = 1, hjust=1))+
  xlab(NULL)
```

The same after recalibration
```{r Fig4B_right,fig.width=3,fig.height=2.4}

m <- subset(d,columns=Condition=="Mock")
m<- FitKinetics(m,type="nlls",time="calibrated_time",compute.residuals=TRUE,return.extra = function(s) setNames(s$residuals$Relative,paste0("Residuals.",s$residuals$Name))[s$residuals$Type=="new"])

df <- reshape2::melt(GetAnalysisTable(m,columns = "Residuals", gene.info = FALSE)[,-1], id.vars=c(),variable.name = "Sample", value.name = "Residual")
df$Sample=plyr::revalue(df$Sample,setNames(sub("kinetics.Mock.Residuals.","",levels(df$Sample)),levels(df$Sample)))
ggplot(df,aes(Sample,Residual,fill=Sample))+
    cowplot::theme_cowplot()+
  geom_hline(yintercept = 0, linetype = 2)+
  geom_boxplot(outlier.size = 0.1)+
  scale_fill_brewer(palette="Reds",guide="none")+
  coord_cartesian(ylim=c(-1,1))+
  scale_y_continuous("Relative residual",labels = scales::percent_format(1))+
  theme(axis.text.x = element_text(angle = 45, vjust = 1, hjust=1))+
  xlab(NULL)
```

Estimate half-lives for all genes before and after recalibration, and compute their log2 fold changes (uncalibrated vs recalibrated). Create boxplots after stratifying for RNA half-lives.

```{r Fig4C,fig.width=3.2,fig.height=2.6}
d <- FitKinetics(d,name="nlls",type="nlls",steady.state=list(Mock=TRUE,SARS=FALSE),return.fields=c("Synthesis","Half-life","f0"))
d <- FitKinetics(d,name="cnlls",type="nlls",steady.state=list(Mock=TRUE,SARS=FALSE),time="calibrated_time",return.fields=c("Synthesis","Half-life","f0"))

df <- GetAnalysisTable(d,columns="Half-life|f0")
df2<-data.frame(
  diff=c(
    log2(df$`nlls.Mock.Half-life`/df$`cnlls.Mock.Half-life`),
    log2(df$`nlls.SARS.Half-life`/df$`cnlls.SARS.Half-life`)
  ),
  `Half-life all`=c(df$`cnlls.Mock.Half-life`,df$`cnlls.SARS.Half-life`),
  Condition=rep(c("Mock","SARS"),each=nrow(df)),
  check.names = FALSE
)
df2$`Half-life`=cut(df2$`Half-life all`,breaks=c(0,1,2,4,8,Inf),labels=c("0-1h","1-2h","2-4h","4-8h",">8h"))
ggplot(df2,aes(`Half-life`,diff,fill=Condition,color=Condition))+
    cowplot::theme_cowplot()+
  geom_boxplot(outlier.size = 0.1)+
  coord_cartesian(ylim=c(-1,1))+
  geom_hline(yintercept = 0)+
  scale_y_continuous(breaks=-1:1)+
  ylab("log FC half-life\nupon calibration")+
  scale_fill_manual(NULL,values=c(Mock='#2b8cbe',`SARS`='#e34a33'))+
  scale_color_manual(NULL,values=c(Mock='#034e7b',`SARS`='#990000'))+
  theme(legend.position = c(0,0),legend.justification = c(-0.1,-0.1),axis.text.x = element_text(angle = 45, vjust = 1, hjust=1))
```

Create the same boxplots for the RNA abundance at t=0 instead of the RNA half-live.

```{r Fig4D,fig.width=3.2,fig.height=2.6}
df2<-data.frame(
  diff=log2(df$`nlls.SARS.f0`/df$`cnlls.SARS.f0`),
  `Half-life all`=c(df$`cnlls.Mock.Half-life`,df$`cnlls.SARS.Half-life`),
  check.names = FALSE
)
df2$`Half-life`=cut(df2$`Half-life all`,breaks=c(0,1,2,4,8,Inf),labels=c("0-1h","1-2h","2-4h","4-8h",">8h"))
ggplot(df2,aes(`Half-life`,diff))+
    cowplot::theme_cowplot()+
  geom_boxplot(color='#990000',fill='#e34a33',outlier.size = 0.1)+
  coord_cartesian(ylim=c(-0.5,0.5))+
  geom_hline(yintercept = 0)+
  scale_y_continuous(breaks=c(-0.5,0,0.5))+
  ylab("log FC RNA abundance\nat t=0\nupon calibration")+
  theme(legend.position = c(0,0),legend.justification = c(-0.1,-0.1),axis.text.x = element_text(angle = 45, vjust = 1, hjust=1))
```
````
