## Supplementary data file 1 for "grandR: a comprehensive package for nucleotide conversion sequencing data analysis": Fig5A-I.Rmd

````
---
title: "Figure 5A-F"
output: 
  html_document:
    dev: "svg"
    keep_md: yes

---

Load the simulation.
To see how this data was generated have a look into the loadGenerate R script.

```{r}
suppressPackageStartupMessages({
    library(grandR)
    library(ggplot2)
    library(patchwork)
})
source("loadGenerate.R")

m=loadGenerate(PerturbationExperiment)
```
Old RNA log fold changes, counter-intuitively do not correspond to log fold changes of the RNA half-life. The hierarchical model is unbiased.

The 4 figures show:

  1. Hierarchical model for synthesis, experiment with perturbed synthesis rates.
  2. Hierarchical model for degradation, experiment with perturbed degradation rates.
  3. New RNA, experiment with perturbed synthesis rates.
  4. Old RNA, experiment with perturbed degradation rates.
  
```{r Fig5A-D, fig.height=2.2, fig.width=2.2, dev='CairoPNG'}
# Plot directly from grandR object
df=GetAnalysisTable(m)
rmsd=mean((log2(df$true_d/df$true_d.perturbed.d)-df$`old.HL vs control.2h.LFC`)^2)
PlotScatter(m, x=log2(true_d/true_d.perturbed.d), y=`old.HL vs control.2h.LFC`,xlim=c(-3,3),ylim=c(-3,3)) + 
  xlab("log FC half-life\n(truth)") + 
  ylab(" \nlog FC old RNA") + 
  annotate('label',x=Inf,y=-Inf,hjust=1.01,vjust=-0.1,label=sprintf("RMSD=%.3f",rmsd),fill='#FFFFFFAA')+
  geom_abline()

rmsd=mean((log2(df$true_d/df$true_d.perturbed.d)-df$`Regulation.HL vs control.2h.HL.log2FC`)^2)
PlotScatter(m, x=log2(true_d/true_d.perturbed.d), y=`Regulation.HL vs control.2h.HL.log2FC`,xlim=c(-3,3),ylim=c(-3,3)) + 
  xlab("log FC half-life\n(truth)") + 
  ylab("log FC half-life\n(MAP)") + 
  annotate('label',x=Inf,y=-Inf,hjust=1.01,vjust=-0.1,label=sprintf("RMSD=%.3f",rmsd),fill='#FFFFFFAA')+
  geom_abline()

rmsd=mean((log2(df$true_s.perturbed.s/df$true_s)-df$`new.s vs control.2h.LFC`)^2)
PlotScatter(m, x=log2(true_s.perturbed.s/true_s), y=`new.s vs control.2h.LFC`,xlim=c(-3,3),ylim=c(-3,3)) + 
  xlab("log FC synthesis\n(truth)") + 
  ylab(" \nlog FC new RNA") + 
  annotate('label',x=Inf,y=-Inf,hjust=1.01,vjust=-0.1,label=sprintf("RMSD=%.3f",rmsd),fill='#FFFFFFAA')+
  geom_abline()

rmsd=mean((log2(df$true_s.perturbed.s/df$true_s)-df$`Regulation.s vs control.2h.s.log2FC`)^2)
PlotScatter(m, x=log2(true_s.perturbed.s/true_s), y=`Regulation.s vs control.2h.s.log2FC`,xlim=c(-3,3),ylim=c(-3,3)) + 
  xlab("log FC synthesis\n(truth)") + 
  ylab("log FC synthesis\n(MAP)") + 
  annotate('label',x=Inf,y=-Inf,hjust=1.01,vjust=-0.1,label=sprintf("RMSD=%.3f",rmsd),fill='#FFFFFFAA')+
  geom_abline()
```

What about new RNA in data with perturbed half-life (or unperturbed as control? For short lived genes, if the degradation rate is changed, also new RNA shows a fold change!

```{r Fig5E, fig.height=3, fig.width=8}
df=GetAnalysisTable(m)
df$`Half-life`=cut(log(2)/df$true_d,breaks=c(0,2,4,6,8,Inf),labels = c("0-2h","2-4h","4-6h","6-8h",">8h"))
types=c("new RNA (perturbed HL)","Synthesis rate (perturbed HL)","new RNA (unperturbed)","Synthesis rate (unperturbed)")
types=factor(types,levels=types[c(1,3,2,4)])
df=rbind(
  data.frame(`Half-life`=df$`Half-life`,`log FC`=df$`new.HL vs control.2h.LFC`,type=types[1],check.names=FALSE),
  data.frame(`Half-life`=df$`Half-life`,`log FC`=df$`Regulation.HL vs control.2h.s.log2FC`-median(df$`Regulation.HL vs control.2h.s.log2FC`),type=types[2],check.names=FALSE),
  data.frame(`Half-life`=df$`Half-life`,`log FC`=df$`new.unperturbed vs control.2h.LFC`,type=types[3],check.names=FALSE),
  data.frame(`Half-life`=df$`Half-life`,`log FC`=df$`Regulation.unperturbed vs control.2h.s.log2FC`-median(df$`Regulation.unperturbed vs control.2h.s.log2FC`),type=types[4],check.names=FALSE)
  )
ggplot(df,aes(`Half-life`,`log FC`,fill=type,color=type))+
    cowplot::theme_cowplot()+
  geom_hline(yintercept = 0)+
  geom_boxplot(outlier.size = 0.3)+
  scale_fill_manual(NULL,values=setNames(c("#d5d5d3","#e2cb49",'white','white'),types))+
  scale_color_manual(NULL,values=setNames(c("#949493","#a09034","#949493","#a09034"),types))+
  coord_cartesian(ylim=c(-1,1))
  
```

If synthesis rate and half-lives are not change, does the hierarchical model recognize this? Compare the distribution of estimated log fold changes (i) unperturbed vs unperturbed, (ii) perturbed synthesis vs unperturbed and (iii) perturbed half-lives vs perturbed synthesis. Do this for all genes with 2h labeling, and for short-lived RNA / 4h labeling.

```{r Fig5F, fig.height=1.7, fig.width=4, warning=FALSE}
types=c("Synthesis rate","Half-life")
exper=c(unperturbed="unperturbed",s="perturbed synthesis",HL="perturbed half-life")

types=factor(types,levels=types)
exper=factor(exper,levels=exper)

tabulate.changes=function(perturbation,timepoint="2h",HL=c(0,Inf)) {
  df=GetAnalysisTable(m)
  df$`Half-life`=log(2)/df$true_d
  df=df[df$`Half-life`>HL[1] & df$`Half-life`<HL[2],]
  rbind(
    data.frame(`log FC`=df[[sprintf("Regulation.%s vs control.%s.s.log2FC",perturbation,timepoint)]],type=types[1],experiment=exper[perturbation],check.names=FALSE),
    data.frame(`log FC`=df[[sprintf("Regulation.%s vs control.%s.HL.log2FC",perturbation,timepoint)]],type=types[2],experiment=exper[perturbation],check.names=FALSE)
  )
}

pl=function(timepoint="2h",HL=c(0,Inf)) {
  df=rbind(tabulate.changes("unperturbed",timepoint,HL),tabulate.changes("s",timepoint,HL),tabulate.changes("HL",timepoint,HL))
  ggplot(df,aes(`log FC`,color=type, linetype=experiment))+
    cowplot::theme_cowplot()+
    stat_ecdf(size=0.7)+
    scale_color_manual(NULL,values=setNames(c("#e2cb49","#3748ae"),types))+
    scale_linetype_manual(NULL,values=c(1,2,3))+
    coord_cartesian(xlim=c(-2,2))+
    xlab("log FC vs control (MAP)")+
    ylab("Cumulative frequency")+
    scale_y_continuous(breaks=c(0,0.5,1),labels=c("0","0.5","1"))+
    scale_x_continuous(breaks=c(-2,0,2),labels=c("-2","0","2"))+
    theme(legend.key.width=unit(0.7,"cm"))+
    guides(linetype = guide_legend(override.aes = list(size = 0.5)))
}

pl()
pl(HL=c(0,2))

pl(timepoint="4h")
pl(timepoint="4h",HL=c(0,2))

```
````
