## Supplementary data file 1 for "grandR: a comprehensive package for nucleotide conversion sequencing data analysis": Fig5G.Rmd

````
---
title: "Figure 5G"
output: 
  html_document:
    dev: "svg"
    keep_md: yes
---

# Comparison of degradation rate changes: full time course versus snapshot

We compare delta between a snapshot from a single time point (2h) versus a
full time course. We will use the real data set as the time course.

```{r}
suppressPackageStartupMessages({
    library(grandR)
    library(ggplot2)
    library(patchwork)
})
source("loadGenerate.R")

sars=loadGenerate(SARS2hRegulation)
```

Make the x and y axis for degradation rate changes:

```{r Fig5G, fig.height=3.5, fig.width=3.5, dev='CairoPNG'}
df=GetAnalysisTable(sars)
PlotScatter(df,x=log2(`nlls.SARS.Half-life`/`nlls.Mock.Half-life`),y=`Regulation.SARS vs Mock.2.HL.log2FC`,xlim=c(-3,3),ylim=c(-3,3),correlation = FormatCorrelation()) + 
  xlab("log FC half-life (full time course)") + 
  ylab("log FC half-life (MAP of 2h)")
```
````
