## Supplementary data file 1 for "grandR: a comprehensive package for nucleotide conversion sequencing data analysis": Fig6.Rmd

````
---
title: "Figure 6"
output: 
  html_document:
    dev: "svg"
    keep_md: yes
---

Load the simulation
To see how this data was generated have a look into the loadGenerate R script.


```{r}
suppressPackageStartupMessages({
    library(grandR)
    library(ggplot2)
    library(patchwork)
})
source("loadGenerate.R")

m=loadGenerate(PerturbationExperiment)
```


# Region of practical equivalence (ROPE) analysis outperforms DESeq2


```{r Fig6A-B,fig.height=2.5, fig.width=5}

tabulate.more.than.X.fold=function(perturbation,timepoint="2h",X=2,HL=c(0,Inf),updown=FALSE) {
  df=GetAnalysisTable(m)
  df$`Half-life`=log(2)/df$true_d
  df=df[df$`Half-life`>HL[1] & df$`Half-life`<HL[2],]
  
  df$truth=if (perturbation=='s') log2(df$true_s.perturbed.s/df$true_s) else log2(df$true_d/df$true_d.perturbed.d)
  df$truth=if (updown) cut(df$truth,breaks=c(-Inf,-log2(X),log2(X),Inf),labels=c("Down","NR","Up")) else cut(abs(df$truth),breaks=c(0,log2(X),Inf),labels=c("NR","Reg"))
  
  s=ifelse(abs(df[,sprintf("Regulation.%s vs control.%s.s.ROPE",perturbation,timepoint)])>0.9,"P>0.9","P<0.9")
  d=ifelse(abs(df[,sprintf("Regulation.%s vs control.%s.HL.ROPE",perturbation,timepoint)])>0.9,"P>0.9","P<0.9")
    
  re=list(table(s,df$truth),table(d,df$truth))
  
  s=ifelse(df[,sprintf("new.%s vs control.%s.Q",perturbation,timepoint)]<0.01,"q<1%","q>1%")
  d=ifelse(df[,sprintf("old.%s vs control.%s.Q",perturbation,timepoint)]<0.01,"q<1%","q>1%")
  
  c(re,list(table(s,df$truth),table(d,df$truth)))
}
format.total=function(tab) setNames(c(sprintf("%.2f%%",100*rowSums(tab/sum(tab))),sprintf("n=%d",rowSums(tab))),rep(rownames(tab),2))
format.updown=function(tab) tab/rep(colSums(tab),each=2)*100

tabulate.rolling=function(perturbation,timepoint="2h",width=200,step=50, HL=c(0,Inf)) {
  df=GetAnalysisTable(m)
  df$`Half-life`=log(2)/df$true_d
  df=df[df$`Half-life`>HL[1] & df$`Half-life`<HL[2],]
  
  df$truth=if (perturbation=='s') log2(df$true_s.perturbed.s/df$true_s) else log2(df$true_d/df$true_d.perturbed.d)
  df=df[order(df$truth),]
  
  
  starts=seq(1,nrow(df)-width+1,by=step)
  rolling=do.call("rbind",lapply(starts,function(st) {
    sub=st:(st+width-1)
    s=abs(df[sub,sprintf("Regulation.%s vs control.%s.s.ROPE",perturbation,timepoint)])>0.9
    d=abs(df[sub,sprintf("Regulation.%s vs control.%s.HL.ROPE",perturbation,timepoint)])>0.9
    re=data.frame(logFC=median(df$truth[sub]),type="ROPE (P>0.9)",regulated=c("Both","Synthesis","Half-life","None"),Frequency=c(sum(s&d),sum(s&!d),sum(!s&d),sum(!s&!d))/length(sub))
    
    s=df[sub,sprintf("new.%s vs control.%s.Q",perturbation,timepoint)]<0.01
    d=df[sub,sprintf("old.%s vs control.%s.Q",perturbation,timepoint)]<0.01
    s[is.na(s)]=FALSE
    d[is.na(d)]=FALSE
    
    re=rbind(re,data.frame(logFC=median(df$truth[sub]),type="DESeq2 (1% FDR)",regulated=c("Both","Synthesis","Half-life","None"),Frequency=c(sum(s&d),sum(s&!d),sum(!s&d),sum(!s&!d))/length(sub)))
    re$regulated=factor(re$regulated,levels=c("None","Synthesis","Half-life","Both"))
    names(re)[1]=if (perturbation=='s') "True log FC (Synthesis)" else "True log FC (Half-life)"
    re
  }))
  rolling$type=factor(rolling$type,levels=c("ROPE (P>0.9)","DESeq2 (1% FDR)"))
  rolling
}
plot.rolling=function(...) {
  rolling=tabulate.rolling(...)
  ggplot(rolling,aes_string(paste0("`",names(rolling)[1],"`"),"Frequency",color="regulated",linetype="type"))+
    cowplot::theme_cowplot()+
    geom_line(size=0.7)+
    coord_cartesian(xlim=c(-2,2))+
    scale_y_continuous(labels=scales::percent_format(1))+
    scale_color_manual("Detected regulation",values=c(None="gray60",Synthesis="#e2cb49",`Half-life`="#3748ae", Both="#b7382b"))+
    scale_linetype_manual(NULL,values=c(1,3))+
    theme(strip.background = element_rect(color="black", fill="white", linetype="solid"))
}


plot.rolling("s")+ggtitle("Perturbed synthesis")
plot.rolling("HL")+ggtitle("Perturbed half-life")


cat("For changed synthesis:\n")
stab=tabulate.more.than.X.fold("s")
cat("detected in degradation and old RNA:\n")
format.total(stab[[2]])
format.total(stab[[4]])

cat("\ndetected in synthesis and new RNA:\n")
stab[[1]]
format.updown(stab[[1]])
stab[[3]]
format.updown(stab[[3]])


cat("\n\nFor changed stability:\n")
stab=tabulate.more.than.X.fold("HL")
stabe=tabulate.more.than.X.fold("HL",updown = TRUE)
cat("detected in synthesis and new RNA:\n")
format.total(stab[[1]])
format.total(stab[[3]])
stab[[3]]
format.updown(stab[[3]])
stabe[[3]]
format.updown(stabe[[3]])

cat("\ndetected in degradation and old RNA:\n")
stab[[2]]
format.updown(stab[[2]])
stab[[4]]
format.updown(stab[[4]])
stabe[[2]]
format.updown(stabe[[2]])
stabe[[4]]
format.updown(stabe[[4]])
        
```

```{r Fig6C, fig.height=1.5, fig.width=2}
plot.rolling("HL",HL=c(0,2))+ggtitle("HL 0-2h (2h)")+guides(color="none",linetype="none")+scale_y_continuous(breaks = c(0,0.5,1),labels=scales::percent_format(1))+ylab(NULL)+xlab(NULL)
plot.rolling("HL",HL=c(0,2),timepoint='4h')+ggtitle("HL 0-2h (4h)")+guides(color="none",linetype="none")+scale_y_continuous(breaks = c(0,0.5,1),labels=scales::percent_format(1))+ylab(NULL)+xlab(NULL)
plot.rolling("HL",HL=c(2,4))+ggtitle("HL 2-4h (2h)")+guides(color="none",linetype="none")+scale_y_continuous(breaks = c(0,0.5,1),labels=scales::percent_format(1))+ylab(NULL)+xlab(NULL)
plot.rolling("HL",HL=c(2,4),timepoint='4h')+ggtitle("HL 2-4h (4h)")+guides(color="none",linetype="none")+scale_y_continuous(breaks = c(0,0.5,1),labels=scales::percent_format(1))+ylab(NULL)+xlab(NULL)
plot.rolling("HL",HL=c(4,8))+ggtitle("HL 4-8h (2h)")+guides(color="none",linetype="none")+scale_y_continuous(breaks = c(0,0.5,1),labels=scales::percent_format(1))+ylab(NULL)+xlab(NULL)
plot.rolling("HL",HL=c(4,8),timepoint='4h')+ggtitle("HL 4-8h (4h)")+guides(color="none",linetype="none")+scale_y_continuous(breaks = c(0,0.5,1),labels=scales::percent_format(1))+ylab(NULL)+xlab(NULL)
plot.rolling("HL",HL=c(8,Inf))+ggtitle("HL >8 (2h)")+guides(color="none",linetype="none")+scale_y_continuous(breaks = c(0,0.5,1),labels=scales::percent_format(1))+ylab(NULL)+xlab(NULL)
plot.rolling("HL",HL=c(8,Inf),timepoint='4h')+ggtitle("HL >8 (4h)")+guides(color="none",linetype="none")+scale_y_continuous(breaks = c(0,0.5,1),labels=scales::percent_format(1))+ylab(NULL)+xlab(NULL)
```
````
