## Supplementary data file 1 for "grandR: a comprehensive package for nucleotide conversion sequencing data analysis": Fig7_S5.Rmd

````
---
title: "Figure 7,S5"
output: 
  html_document:
    dev: "svg"
    keep_md: yes
---

Read BANP data.
To see how this data was generated have a look into the loadGenerate R script.


```{r}
suppressPackageStartupMessages({
    library(grandR)
    library(ggplot2)
    library(patchwork)
})
source("loadGenerate.R")

d=loadGenerate(BanpRegulation)
colors=setNames(c("black",rev(RColorBrewer::brewer.pal(8,"Greens"))[c(2:5,8)]),c("0h","1h","2h","4h","6h","20h"))
```

Distribution of Half-lives estimated separately for each time point after recalibration.
```{r FigS5A, fig.height=2.5, fig.width=4}
df=GetAnalysisTable(d,analyses = "^cal.MAP",columns = "^HL$",gene.info = FALSE)
names(df)=names(colors)
df=reshape2::melt(df,id.vars = c(),variable.name = "timepoint", value.name = "Half-life [h]")
ggplot(df,aes(timepoint,`Half-life [h]`,fill=timepoint))+
    cowplot::theme_cowplot()+
  geom_violin(color='black',size=0.1)+
  scale_fill_manual(NULL,values=colors,guide="none")+
  scale_y_log10(breaks=c(1,5,24))+
  coord_cartesian(ylim=c(0.5,35))+
  xlab("Time after depletion of BANP")
```

Distribution of synthesis rates estimated separately for each time point after recalibration.
```{r FigS5B, fig.height=2.5, fig.width=4}
df=GetAnalysisTable(d,analyses = "^cal.MAP",columns = "^s$",gene.info = FALSE)
names(df)=names(colors)
df=reshape2::melt(df,id.vars = c(),variable.name = "timepoint", value.name = "syn")
ggplot(df,aes(timepoint,`syn`,fill=timepoint))+
    cowplot::theme_cowplot()+
  geom_violin(color='black',size=0.1)+
  scale_fill_manual(NULL,values=colors,guide="none")+
  scale_y_log10("Synthesis rate\n[normalized expression/h]",breaks=c(10,100,1000,10000))+
  coord_cartesian(ylim=c(10,1E4))+
  xlab("Time after depletion of BANP")
```

Compare the distribution of synthesis rate log2 fold changes for target and non-targets for each time point:

```{r Fig7A, fig.height=2, fig.width=5.4, warning=FALSE}
dfg=GetAnalysisTable(d)[,c("Symbol","BANP-Target","Regulation.1h.s.log2FC","Regulation.2h.s.log2FC","Regulation.4h.s.log2FC","Regulation.6h.s.log2FC","Regulation.20h.s.log2FC")]
names(dfg)[-1:-2]=names(colors[-1])
norm=apply(as.matrix(dfg[,-1:-2]),2,median,na.rm=T)
dfg[,-1:-2]=as.data.frame(t(t(as.matrix(dfg[,-1:-2]))-norm))
dfg$`BANP-Target`=factor(dfg$`BANP-Target`,levels=c("no","yes"))

dfg=reshape2::melt(dfg,id.vars = c("Symbol","BANP-Target"),variable.name = "timepoint", value.name = "logFC")
ggplot(dfg,aes(timepoint,logFC,fill=timepoint,linetype=`BANP-Target`,alpha=`BANP-Target`))+
    cowplot::theme_cowplot()+
  geom_boxplot(outlier.shape = NA,position = position_dodge(width = 0.5))+
  geom_hline(yintercept = 0)+
  scale_fill_manual(NULL,values=colors,guide="none")+
  coord_cartesian(ylim=c(-2,1))+
  scale_y_continuous(bquote(log[2]~"FC synthesis rate vs 0h"),breaks=c(-2,-1,0,1))+
  scale_alpha_manual(NULL,values=c(0.3,1),guide="none")+
  scale_linetype_manual(,values=c(3,1))
```

Show ridgeplots for the posterior distributions of synthesis rate fold changes for each time point for 3 example genes.
```{r Fig7B,fig.height=2, fig.width=4, warning=FALSE}
contrasts=GetContrasts(d,contrast=c("Experimental.time.original","0h"),group = "Genotype")
names(contrasts)=gsub(" vs .*","",names(contrasts))
reference.columns=FindReferences(d,reference.function=function(s) Experimental.time==max(c(0,Experimental.time[Experimental.time<s$Experimental.time])),group="Genotype")

pdf=function(gene) {
  set.seed(1337)
  ref=rep(FitKineticsGeneSnapshot(d,gene,contrasts[[1]]==-1,reference.columns = reference.columns,return.samples = TRUE,N = 10000,time.labeling = "calibrated_time",correct.labeling = TRUE)$samples$s,length.out=10000)
  df=do.call("rbind",lapply(names(contrasts),function(name) data.frame(logFC=log2(rep(FitKineticsGeneSnapshot(d,gene,contrasts[[name]]==1,reference.columns = reference.columns,return.samples = TRUE,N=10000,time.labeling = "calibrated_time",correct.labeling = TRUE)$samples$s,length.out=10000)/ref)-norm[name],time=name)))
  df$time=factor(df$time,levels=rev(names(contrasts)))
  df$gene=gene
  df
}
#df=rbind(pdf("Tubgcp5"),pdf("Nsun3"),pdf("Wdr77"),pdf("Taf1d"),pdf("Herc1"),pdf("Churc1"))
genes=c("Taf1d","Herc1","Tubgcp5")
df=do.call("rbind",lapply(genes,pdf))

df=rbind(df,data.frame(logFC=0,time=rep(levels(df$time),each=length(unique(df$gene))),gene=rep(unique(df$gene),length(unique(df$time)))))

pp=function(gene) ggplot(df[df$gene==gene,],aes(logFC,time,fill=time))+
    cowplot::theme_cowplot()+
  ggridges::geom_density_ridges2(size=0.3)+
  ggtitle(gene)+
  geom_vline(xintercept = 0,linetype=3)+
  scale_fill_manual(NULL,values=colors,guide="none")+
  ylab(NULL)+
  xlab(bquote(log[2]~"FC posterior (synthesis rate vs 0h)"))
  


for (gene in genes) print(pp(gene))

```
````
