## Supplementary data file 1 for "grandR: a comprehensive package for nucleotide conversion sequencing data analysis": FigS4.Rmd

````
---
title: "Figure S4A-B"
output: 
  html_document:
    dev: "svg"
    keep_md: yes
---

Load the simulated recalibration data.
To see how this data was generated have a look into the loadGenerate R script.

```{r}
suppressPackageStartupMessages({
    library(grandR)
    library(ggplot2)
    library(patchwork)
})
source("loadGenerate.R")

m=loadGenerate(CalibrateSimulation)

```

Show the correction of effective labeling times (more precisely: arrows pointing from the log2 fold change of the nominal vs. the effective labeling time towards the log2 fold change of the recalibrated vs. the effective labeling time.

```{r Fig S4A,fig.width=4,fig.height=3}
df=Coldata(m)
df$`Uneffective labeling`=paste0(gsub(".*_","",df$Condition),"min")
df$`Steady state`=factor(ifelse(substr(df$Condition,1,1)=='n',"no","yes"),levels=c("yes","no"))
df=droplevels(df[df$duration.4sU<8 & df$duration.4sU>0,])
df$x=as.integer(as.factor(df$duration.4sU))
df$x=df$x+as.integer(as.factor(df$Replicate))*0.1-0.2

ggplot(df,aes(x,log2(duration.4sU/duration.4sU.effective),xend=x,yend=log2(calibrated_time/duration.4sU.effective),color=Replicate))+
  geom_hline(yintercept = 0)+
  geom_segment(arrow = arrow(length = unit(0.1, "cm"))) +
  ylab(bquote(log[2]~"FC labeling time / truth"))+
  facet_grid(`Steady state`~`Uneffective labeling`)+
  coord_cartesian(ylim=c(-1,1.8))+
  scale_color_brewer(NULL,palette="Dark2",guide="none")+
  theme(strip.background = element_rect(color='black',size=0.3,fill = 'white'),panel.background = element_rect(color='black',size=0.3,fill = 'white'))+
  scale_x_continuous("timepoint",breaks=1:3,labels = c("1h","2h","4h"))+
  scale_y_continuous(breaks=c(-1,0,1))

```

Show the effect of recalibration on estimated half-lives.

```{r Fig S4B, fig.width=4,fig.height=3}
tab = GetAnalysisTable(m,columns=c('Half-life$'),gene.info = FALSE)

tab=as.data.frame(lapply(tab,function(hl) log2(hl*GeneInfo(m)$true_d/log(2))),check.names=FALSE)
tab=reshape2::melt(tab,id.vars = c(),variable.name = "Setting",value.name = "logFC")
tab=cbind(tab,MakeColdata(gsub("_",".",tab$Setting),design=c("Mode","Steady state","Uneffective labeling","HL"),rownames = FALSE))
tab$`Steady state`=factor(ifelse(tab$`Steady state`=="ss","yes","no"),levels=c("yes","no"))
tab$`Uneffective labeling`=paste0(tab$`Uneffective labeling`,"min")

ggplot(tab, aes(Mode,y=logFC,color=Mode,fill=Mode)) + 
  geom_boxplot(outlier.shape = NA,size=0.2)+
  coord_cartesian(ylim = c(-0.25,1))+
  theme(axis.text.x = element_text(angle=90, hjust = 1, vjust = 0.5)) + 
  xlab(NULL) + 
  ylab(bquote(log[2]~FC~"estimated HL / truth")) +
  facet_grid(`Steady state`~`Uneffective labeling`,scales = "free_y")+
  scale_fill_manual(NULL,values=c(uncalibrated="#DE2D26",calibrated="#3182BD",truth="white"),guide="none")+
  scale_color_manual(NULL,values=c(uncalibrated="#A50F15",calibrated="#08519C",truth="black"),guide="none")+
  theme(strip.background = element_rect(color='black',size=0.3,fill = 'white'),panel.background = element_rect(color='black',size=0.3,fill = 'white'))
  

```
````
