## Supplementary data file 1 for "grandR: a comprehensive package for nucleotide conversion sequencing data analysis": loadGenerate.R

```
loadGenerateAll=function() {
  for (rmd in list.files(".",".*\\.Rmd$")) {
    for (l in readLines(rmd)) {
      if (grepl("loadGenerate\\(.*\\)",l)) {
        FUN=gsub(".*loadGenerate\\((.*)\\)","\\1",l)
        loadGenerate(FUN,as.char = TRUE)
      }
    }
  }
}

loadGenerate=function(fun,as.char = FALSE) {
  if (!as.char) fun = as.character(substitute(fun))
  dir.create("../data",showWarnings = FALSE)
  file=paste0("../data/",fun,".rds")
	if(file.exists(file)) {
		cat(sprintf("Loaded file %s for %s\n",file,fun))
		readRDS(file)
	} else {
		cat(sprintf("Generating file %s for %s... this will take a while!\n",file,fun))
		re=rlang::exec(fun)
		saveRDS(re, file)
		re
	}
}

ProgressiveLabelingTimecourse=function() {

	sars <- ReadGRAND("https://zenodo.org/record/5834034/files/sars.tsv.gz",design=c(Design$Condition,Design$dur.4sU,Design$Replicate))

	sars = subset(sars, columns=Condition=="Mock")
	sars = FilterGenes(sars)
	sars = Normalize(sars)

	SetParallel()
	sars = FitKinetics(sars, type="nlls")

	seed=1337
	noss=SimulateTimeCourse("noss",GeneInfo(sars),
		                  s = GetAnalysisTable(sars)$kinetics.Mock.Synthesis,
		                  d = log(2)/GetAnalysisTable(sars)$`kinetics.Mock.Half-life`,
		                  f0 = rowMeans(GetTable(sars,type="norm")) *2^rnorm(nrow(sars),0,1),
		                  dispersion=estimate.dispersion(GetTable(sars,type="count")),
		                  s.variation = 1.05,
		                  d.variation = 1.05,
		                  timepoints = c(0, 0, 0, 1, 1, 1, 2, 2, 2, 4, 4, 4, 8, 8, 8),
		                  beta.approx = TRUE,
		                  conversion.reads = TRUE,
		                  num.reads = 2E7,
		                  seed=seed)
	ss=SimulateTimeCourse("ss",GeneInfo(sars),
		                s = GetAnalysisTable(sars)$kinetics.Mock.Synthesis,
		                d = log(2)/GetAnalysisTable(sars)$`kinetics.Mock.Half-life`,
		                dispersion=estimate.dispersion(GetTable(sars,type="count")),
		                s.variation = 1.05,
		                d.variation = 1.05,
		                timepoints = c(0, 0, 0, 1, 1, 1, 2, 2, 2, 4, 4, 4, 8, 8, 8),
		                beta.approx = TRUE,
		                conversion.reads = TRUE,
		                num.reads = 2E7,
		                seed=seed)
	m=merge(ss=ss,noss=noss)

	spike=(abs(log2(GeneInfo(m)$true_f0.noss/rowMeans(GetTable(sars,type="norm"))))<0.0045)
	m = Normalize(m,genes=spike)
	m
}

ProgressiveLabelingTimecourseAllModelsFitted=function() {
  m=loadGenerate(ProgressiveLabelingTimecourse)

	SetParallel()
	
	extra=function(s) c(`Synthesis.lower`=unname(s$conf.lower["Synthesis"]),`Synthesis.upper`=unname(s$conf.upper["Synthesis"]),`Half-life.lower`=unname(s$conf.lower["Half-life"]),`Half-life.upper`=unname(s$conf.upper["Half-life"]))
	m = FitKinetics(m,name = "nlls",type = "nlls", steady.state = c(ss=TRUE,noss=FALSE), return.extra = extra)
	m = FitKinetics(m,name = "ntr",type = "ntr", return.extra = extra, exact.ci=FALSE)
	m = FitKinetics(m,name = "entr",type = "ntr", return.extra = extra, exact.ci=TRUE)
	m = FitKinetics(m,name = "lm",type = "lm", return.extra = extra)
	m = FitKineticsPulseR(m, name = "pulseR")

	m
}

ProgressiveLabelingSettings=function() {

  sars <- ReadGRAND("https://zenodo.org/record/5834034/files/sars.tsv.gz",design=c(Design$Condition,Design$dur.4sU,Design$Replicate))

  sars = subset(sars, columns=Condition=="Mock")
  sars = FilterGenes(sars)
  sars = Normalize(sars)

  SetParallel()
  sars = FitKinetics(sars, type="nlls")

  seed=1337
  m=SimulateTimeCourse("ss",GeneInfo(sars),
                       s = GetAnalysisTable(sars)$kinetics.Mock.Synthesis,
                       d = log(2)/GetAnalysisTable(sars)$`kinetics.Mock.Half-life`,
                       dispersion=estimate.dispersion(GetTable(sars,type="count")),
                       s.variation = 1.05,
                       d.variation = 1.05,
                       timepoints = rep(c(0,1,2,4,8),each=4),
                       num.reads = 2E7,
                       seed=seed)

  m = Normalize(m)
  settings=c(
    "2x 0+1h","2x 0+2h","2x 0+4h","2x 0+8h",
    "4x 0+1h","4x 0+2h","4x 0+4h","4x 0+8h",
    "2x 0+1+2h","2x 0+2+4h","2x 0+4+8h",
    "4x 0+1+2h","4x 0+2+4h","4x 0+4+8h",
    "1x 0+1+2+4h","1x 0+1+2+8h","1x 0+1+4+8h","1x 0+2+4+8h",
    "3x 0+1+2+4h","3x 0+1+2+8h","3x 0+1+4+8h","3x 0+2+4+8h",
    "1x 0+1+2+4+8h","2x 0+1+2+4+8h","3x 0+1+2+4+8h"
  )
  for (s in settings) {
    cat(sprintf("Fitting %s...\n",s))
    reps=as.integer(substr(s,1,1))
    timepoints=as.integer(strsplit(substr(s,4,nchar(s)-1),"\\+")[[1]])
    sub=subset(m,Replicate %in% LETTERS[1:reps] & duration.4sU %in% timepoints)
    sub=DropAnalysis(sub)
    atab=GetAnalysisTable(FitKinetics(sub,name = s,type = "nlls"),gene.info = FALSE,prefix.by.analysis = FALSE)
    m=AddAnalysis(m,s,atab)
  }

  m
}


ProgressiveLabelingDepth=function() {

  sars <- ReadGRAND("https://zenodo.org/record/5834034/files/sars.tsv.gz",design=c(Design$Condition,Design$dur.4sU,Design$Replicate))

  sars = subset(sars, columns=Condition=="Mock")
  sars = FilterGenes(sars)
  sars = Normalize(sars)

  SetParallel()
  sars = FitKinetics(sars, type="nlls")

  seed=1337

  reads=c(5,10,20,50)
  list=lapply(reads,function(r) SimulateTimeCourse(paste0(r,"M"),GeneInfo(sars),
                       s = GetAnalysisTable(sars)$kinetics.Mock.Synthesis,
                       d = log(2)/GetAnalysisTable(sars)$`kinetics.Mock.Half-life`,
                       dispersion=estimate.dispersion(GetTable(sars,type="count")),
                       s.variation = 1.05,
                       d.variation = 1.05,
                       timepoints = rep(c(0,1,2,4,8),each=4),
                       num.reads = r*1E6,
                       seed=seed))
  names(list)=paste0(reads,"M")
  list=lapply(names(list),function(name) {
    m = Normalize(list[[name]])
    for (reps in 1:4) {
      cat(sprintf("Fitting %s %d reps...\n",name,reps))
      sub=subset(m,Replicate %in% LETTERS[1:reps])
      sub=DropAnalysis(sub)
      nn=paste0(name,".",reps,"R")
      atab=GetAnalysisTable(FitKinetics(sub,name = nn,type = "nlls"),gene.info = FALSE,prefix.by.analysis = FALSE)
      m=AddAnalysis(m,nn,atab)
    }
    m
  })
  names(list)=paste0(reads,"M")
  m=merge.grandR(list=list)

  m
}

CalibrateSARS=function() {

  d <- ReadGRAND("https://zenodo.org/record/5834034/files/sars.tsv.gz",design=c(Design$Condition,Design$dur.4sU,Design$Replicate))
  d <- FilterGenes(d)
  d <- Normalize(d)
  SetParallel()
  d <- CalibrateEffectiveLabelingTimeKineticFit(d,steady.state=list(Mock=TRUE,SARS=FALSE),verbose = TRUE)
  d
}


CalibrateSimulation=function() {
  sars <- ReadGRAND("https://zenodo.org/record/5834034/files/sars.tsv.gz",design=c(Design$Condition,Design$dur.4sU,Design$Replicate))

  sars = subset(sars, columns=Condition=="Mock")
  sars = FilterGenes(sars)
  sars = Normalize(sars)

  SetParallel()
  sars = FitKinetics(sars, type="nlls")

  create=function(uneff.labeling) {
    seed=1337
    noss=SimulateTimeCourse(paste0("noss_",uneff.labeling),GeneInfo(sars),
                            s = GetAnalysisTable(sars)$kinetics.Mock.Synthesis,
                            d = log(2)/GetAnalysisTable(sars)$`kinetics.Mock.Half-life`,
                            f0 = rowMeans(GetTable(sars,type="norm")) *2^rnorm(nrow(sars),0,1),
                            dispersion=estimate.dispersion(GetTable(sars,type="count")),
                            s.variation = 1.05,
                            d.variation = 1.05,
                            timepoints = rep(c(0,1-uneff.labeling/60,2-uneff.labeling/60,4-uneff.labeling/60,8),each=3),
                            num.reads = 2E7,
                            seed=seed)
    Coldata(noss,"duration.4sU.effective")=Coldata(noss,"duration.4sU")
    Coldata(noss,"duration.4sU")=rep(c(0,1,2,4,8),each=3)
    Coldata(noss,"duration.4sU.original")=NULL
    ss=SimulateTimeCourse(paste0("ss_",uneff.labeling),GeneInfo(sars),
                          s = GetAnalysisTable(sars)$kinetics.Mock.Synthesis,
                          d = log(2)/GetAnalysisTable(sars)$`kinetics.Mock.Half-life`,
                          dispersion=estimate.dispersion(GetTable(sars,type="count")),
                          s.variation = 1.05,
                          d.variation = 1.05,
                          timepoints = rep(c(0,1-uneff.labeling/60,2-uneff.labeling/60,4-uneff.labeling/60,8),each=3),
                          num.reads = 2E7,
                          seed=seed)
    Coldata(ss,"duration.4sU.effective")=Coldata(ss,"duration.4sU")
    Coldata(ss,"duration.4sU")=rep(c(0,1,2,4,8),each=3)
    Coldata(ss,"duration.4sU.original")=NULL
    setNames(list(ss,noss),paste0(c("ss_","noss_"),uneff.labeling))
  }
  m.list=c(create(20),create(30),create(40))
  m=merge.grandR(list=m.list)

  spike=(abs(log2(GeneInfo(m)$true_f0.noss_20/rowMeans(GetTable(sars,type="norm"))))<0.0045)
  m = Normalize(m,genes=spike)


  stst=setNames(as.list(!grepl("noss",names(m.list))),names(m.list))
  m=CalibrateEffectiveLabelingTimeKineticFit(m,steady.state=stst,verbose=TRUE)

  m=FitKinetics(m,name="uncalibrated",type="nlls",time="duration.4sU",steady.state = stst)
  m=FitKinetics(m,name="calibrated",type="nlls",time="calibrated_time",steady.state = stst)
  m=FitKinetics(m,name="truth",type="nlls",time="duration.4sU.effective",steady.state = stst)

  m
}


PerturbationExperiment=function() {
  sars <- ReadGRAND("https://zenodo.org/record/5834034/files/sars.tsv.gz",design=c(Design$Condition,Design$dur.4sU,Design$Replicate))

  sars = subset(sars, columns=Condition=="Mock")
  sars = FilterGenes(sars)
  sars = Normalize(sars)

  SetParallel()
  sars = FitKinetics(sars, type="nlls")

  seed=1337
  f0=GetAnalysisTable(sars)$kinetics.Mock.Synthesis/log(2)*GetAnalysisTable(sars)$`kinetics.Mock.Half-life`
  control=SimulateTimeCourse("control",GeneInfo(sars),
                             s = GetAnalysisTable(sars)$kinetics.Mock.Synthesis,
                             d = log(2)/GetAnalysisTable(sars)$`kinetics.Mock.Half-life`,
                             dispersion=estimate.dispersion(GetTable(sars,type="count")),
                             f0 = f0,
                             s.variation = 1.05,
                             d.variation = 1.05,
                             timepoints = c(1,1,1,2,2,2,4,4,4,8,8,8),
                             beta.approx = TRUE,
                             num.reads=2E7,
                             seed=seed)
  unperturbed=SimulateTimeCourse("unperturbed",GeneInfo(sars),
                                 s = GetAnalysisTable(sars)$kinetics.Mock.Synthesis,
                                 d = log(2)/GetAnalysisTable(sars)$`kinetics.Mock.Half-life`,
                                 dispersion=estimate.dispersion(GetTable(sars,type="count")),
                                 f0 = f0,
                                 s.variation = 1.05,
                                 d.variation = 1.05,
                                 timepoints = c(1,1,1,2,2,2,4,4,4,8,8,8),
                                 beta.approx = TRUE,
                                 num.reads=2E7,
                                 seed=seed*13)
  perturbed.d=SimulateTimeCourse("HL",GeneInfo(sars),
                                 s = GetAnalysisTable(sars)$kinetics.Mock.Synthesis,
                                 d = log(2)/GetAnalysisTable(sars)$`kinetics.Mock.Half-life` *2^rnorm(nrow(sars),0,1) ,
                                 dispersion=estimate.dispersion(GetTable(sars,type="count")),
                                 f0 = f0,
                                 s.variation = 1.05,
                                 d.variation = 1.05,
                                 timepoints = c(1,1,1,2,2,2,4,4,4,8,8,8),
                                 beta.approx = TRUE,
                                 num.reads=2E7,
                                 seed=seed*13*13)
  perturbed.s=SimulateTimeCourse("s",GeneInfo(sars),
                                 s = GetAnalysisTable(sars)$kinetics.Mock.Synthesis *2^rnorm(nrow(sars),0,1) ,
                                 d = log(2)/GetAnalysisTable(sars)$`kinetics.Mock.Half-life`,
                                 dispersion=estimate.dispersion(GetTable(sars,type="count")),
                                 f0 = f0,
                                 s.variation = 1.05,
                                 d.variation = 1.05,
                                 timepoints = c(1,1,1,2,2,2,4,4,4,8,8,8),
                                 beta.approx = TRUE,
                                 num.reads=2E7,
                                 seed=seed*13*13*13)

  m=merge(control=control,unperturbed=unperturbed,perturbed.d=perturbed.d,perturbed.s=perturbed.s)

  m=Normalize(m)

  SetParallel()
  m=PairwiseDESeq2(m,"total",GetContrasts(m,contrast = c("Condition","control"),group = "duration.4sU.original"),mode="total",verbose=T,separate = T)
  m=LFC(m,"total",GetContrasts(m,contrast = c("Condition","control"),group = "duration.4sU.original"),mode="total")
  m=PairwiseDESeq2(m,"new",GetContrasts(m,contrast = c("Condition","control"),group = "duration.4sU.original"),mode="new",verbose=T,separate = T)
  m=LFC(m,"new",GetContrasts(m,contrast = c("Condition","control"),group = "duration.4sU.original"),mode="new")
  m=PairwiseDESeq2(m,"old",GetContrasts(m,contrast = c("Condition","control"),group = "duration.4sU.original"),mode="old",verbose=T,separate = T)
  m=LFC(m,"old",GetContrasts(m,contrast = c("Condition","control"),group = "duration.4sU.original"),mode="old")


  m=EstimateRegulation(m,"Regulation",
                       contrasts = GetContrasts(m,contrast = c("Condition","control"),group = "duration.4sU.original"),
                       reference.columns=FindReferences(m,Condition=="control",group="duration.4sU.original"),
                       verbose=TRUE,
                       sample.f0.in.ss = TRUE,N=10)
  m
}

# TODO: rerun
SARS2hRegulation=function() {
  sars=loadGenerate(CalibrateSARS)

  SetParallel()
  sars <- FitKinetics(sars, "nlls", steady.state=c(Mock=TRUE,SARS=FALSE),time="calibrated_time")
  sars <- EstimateRegulation(sars,
                             contrasts = GetContrasts(sars, contrast = c("Condition","Mock"),group="duration.4sU",columns=Coldata(sars)$duration.4sU==2),
                             reference.columns = FindReferences(sars, duration.4sU==0,group="Condition"),
                             time.labeling = "calibrated_time",
                             correct.labeling=TRUE,
                             verbose = TRUE)
  sars
}

BanpRegulation=function() {

  d=ReadGRAND("https://zenodo.org/record/6976391/files/BANP.tsv.gz",design=c("Condition","Experimental.time","Genotype",Design$dur.4sU,Design$has.4sU,"Replicate"))
  d=FilterGenes(d)
  d=Normalize(d)
  tar=readLines("https://zenodo.org/record/6976391/files/targets.genes")
  GeneInfo(d,"BANP-Target")<-factor(ifelse(Genes(d) %in% tar,"yes","no"),levels=c("yes","no"))
  d=subset(d,columns = Genotype=="dTag")

  contrasts=GetContrasts(d,contrast=c("Experimental.time.original","0h"),name.format="$A")
  reference.columns=FindReferences(d,reference.function=function(s) Experimental.time==max(c(0,Experimental.time[Experimental.time<s$Experimental.time])),group="Genotype")


  SetParallel()

  d=ComputeSteadyStateHalfLives(d,name="HL",columns = Experimental.time==0)
  GeneInfo(d,"HL simple") <- rowMeans(GetAnalysisTable(d,analyses = "HL",gene.info = FALSE))
  d=CalibrateEffectiveLabelingTimeMatchHalflives(d,reference.halflives = GeneInfo(d,"HL simple"),reference.columns = reference.columns)

  Condition(d) <- "Experimental.time.original"
  d <- FitKineticsSnapshot(d,name.prefix = "uncal.MAP",reference.columns=reference.columns,time.experiment = "Experimental.time",N=0)
  d <- FitKineticsSnapshot(d,name.prefix = "cal.MAP",reference.columns=reference.columns,time.experiment = "Experimental.time",N=0,time.labeling = "calibrated_time",correct.labeling = TRUE)

  d=EstimateRegulation(d,"Regulation",contrasts=contrasts,reference.columns=reference.columns,time.labeling = "calibrated_time",verbose=T,time.experiment = "Experimental.time",correct.labeling=TRUE,N=0)

  d
}
```
